# Suppression of astrocyte BMP signaling improves fragile X syndrome molecular signatures and functional deficits

**DOI:** 10.1101/2024.06.19.599752

**Authors:** James Deng, Lara Labarta-Bajo, Ashley N. Brandebura, Samuel B. Kahn, Antonio F. M. Pinto, Jolene K. Diedrich, Nicola J. Allen

## Abstract

Fragile X syndrome (FXS) is a monogenic neurodevelopmental disorder with manifestations spanning molecular, neuroanatomical, and behavioral changes. Astrocytes contribute to FXS pathogenesis and show hundreds of dysregulated genes and proteins; targeting upstream pathways mediating astrocyte changes in FXS could therefore be a point of intervention. To address this, we focused on the bone morphogenetic protein (BMP) pathway, which is upregulated in FXS astrocytes. We generated a conditional KO (cKO) of Smad4 in astrocytes to suppress BMP signaling, and found this lessens audiogenic seizure severity in FXS mice. To ask how this occurs on a molecular level, we performed *in vivo* transcriptomic and proteomic profiling of cortical astrocytes, finding upregulation of metabolic pathways, and downregulation of secretory machinery and secreted proteins in FXS astrocytes, with these alterations no longer present when BMP signaling is suppressed. Functionally, astrocyte Smad4 cKO restores deficits in inhibitory synapses present in FXS auditory cortex. Thus, astrocytes contribute to FXS molecular and functional phenotypes, and targeting astrocytes can mitigate FXS symptoms.

## Main Text

Fragile X syndrome (FXS) is the most common single-gene cause and most common inherited form of intellectual disability and autism spectrum disorder (ASD).^1,2^ Loss of fragile X messenger ribonucleoprotein (FMRP, gene = *fragile X messenger ribonucleoprotein 1*, *Fmr1*) causes FXS, an X-linked disorder with manifestations including immature dendritic spines on neurons, hyperarousal and hypersensitivity across sensory modalities, and seizures.^1,3,4^ Research in FXS and with Fmr1 knock-out (KO) mice, the most common animal model of FXS,^5^ has predominantly focused on neuronal dysfunction. However, emerging research suggests that astrocytes, an abundant glial cell type in the CNS, also play a role.^6–8^ During development, astrocytes secrete temporally regulated signals to promote dendrite growth and synapse formation, maturation, and function.^9^ Astrocyte-specific Fmr1 KO and re-expression studies highlight a role for astrocytes in some Fmr1 KO cortical excitability and behavioral phenotypes.^10^

Astrocytes are dysregulated at a molecular level in Fmr1 KO. Prior work has identified hundreds of genes and proteins altered in FXS astrocytes both *in vitro* and *in vivo*.^11,12^ Moreover, cell culture experiments show bone morphogenetic protein (BMP) signaling is an upstream regulatory pathway that drives FXS-like changes to astrocyte secreted proteins,^12^ and that astrocyte-secreted factors can mediate FXS neuronal dysfunction.^12,13^ Additionally, recent work demonstrated that if all cells are heterozygous for the BMP receptor Bmpr2, this can partially rescue FXS dendritic spine defects *in vivo*,^14^ and that Fmr1 KO mice have elevated astrocyte canonical BMP signaling *in vivo* in early development.^12^ These experiments suggest that BMP signaling may be a mediator of FXS pathology, however, the precise role of astrocyte BMP signaling in Fmr1 KO has not been studied *in vivo*.

BMPs are secreted members of the transforming growth factor β (TGFβ) superfamily and bind to heterotetramers of serine-threonine kinase receptors BMPR1A, BMPR1B, and BMPR2. BMP signaling is mediated canonically by SMAD-dependent pathways leading to transcriptional changes through a complex including SMAD4 (**Fig. 1A**).^15,16^ BMPs can also signal through noncanonical pathways such as via interaction of BMPR2 with LIM Kinase 1.^14,16^ BMP signaling is involved in embryonic neurodevelopmental processes^16^ as well as synaptogenesis^17–19^ and dendritic outgrowth.^20–23^ BMP signaling also acts on astrocytes: BMPs promote differentiation of glial progenitors into astrocytes,^24^ and BMPs lead to maturation of astrocytes *in vitro* with involvement of canonical signaling.^25^ However, BMP signaling is highly developmental state- and context-dependent,^26^ and the role of BMP signaling in astrocytes during postnatal development and in disease remains poorly understood. We hypothesize that astrocyte BMP signaling contributes to FXS deficits, and ask if modulation of this upstream pathway in astrocytes *in vivo* can reverse FXS astrocyte molecular signatures and improve FXS behavioral deficits.

**Figure 1.**
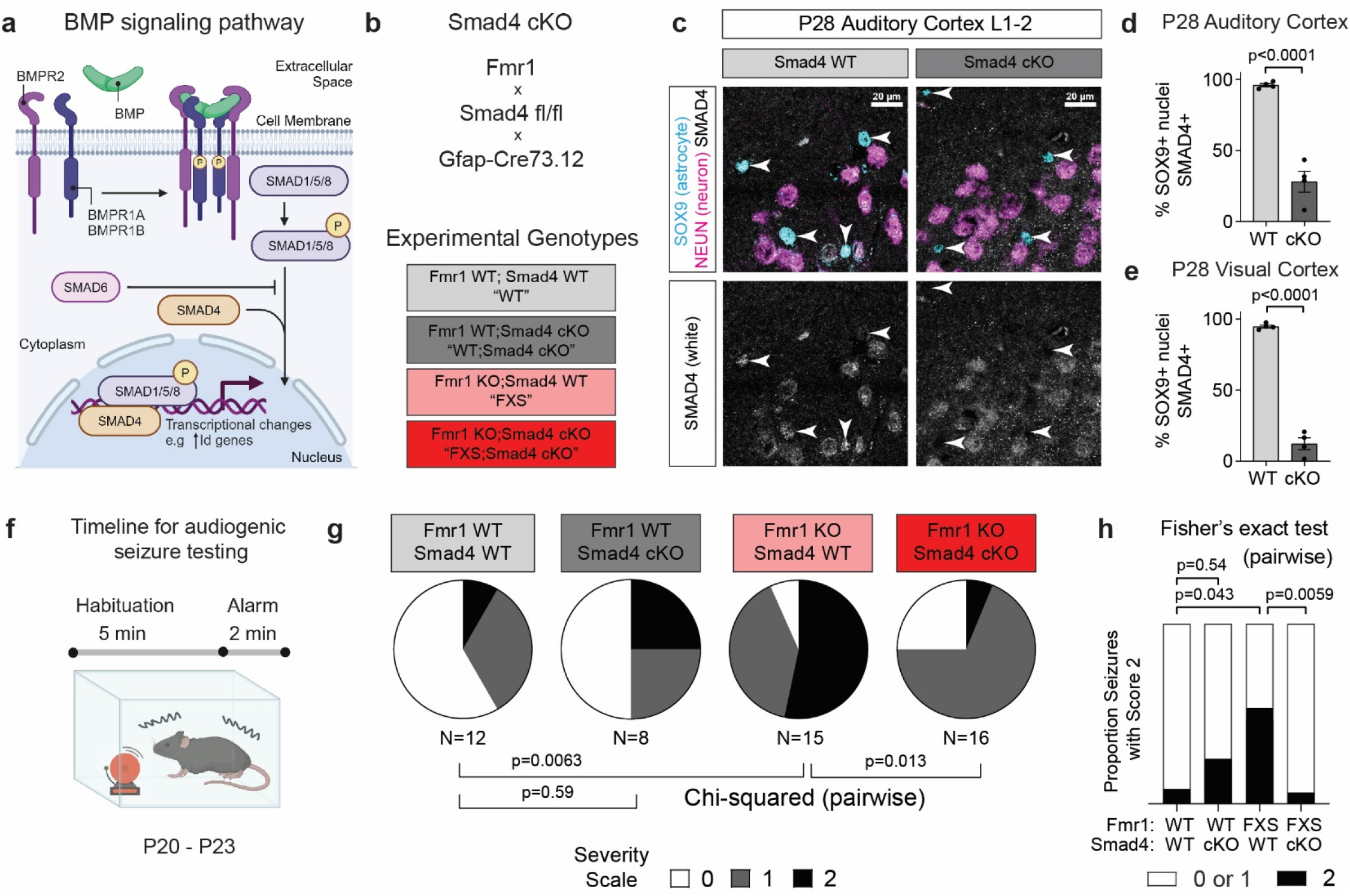
Smad4 cKO suppresses astrocyte BMP signaling and reduces Fmr1 KO audiogenic seizure severity. **a.** Schematic of the canonical BMP pathway. BMPs bind to BMPR, leading to phosphorylation of SMAD1/5/8; pSMAD forms a complex with SMAD4, which translocates to the nucleus and regulates transcription. **b**. Above, outline of mouse genetics strategy to achieve Smad4 conditional knock-out in astrocytes; below, experimental genotype groups. **c**. Smad4 cKO deletes Smad4 from astrocytes. Example images of L1 to L2 auditory cortex, immunohistochemistry for SMAD4, SOX9 (astrocyte nuclei), and NEUN (neuronal nuclei) at P28. White arrowheads indicate locations of SOX9+ astrocyte nuclei. **d**. Quantification across P28 auditory cortex, percentage of SOX9+ nuclei expressing SMAD4 higher than the parenchymal background. N = 4 mice/condition. Statistics by t-test. **e**. P28 visual cortex data, analyzed as in **d**. N = 4 mice/condition. Statistics by t-test. **f**. Timeline for audiogenic seizure testing at P20-23. **g**. Quantification of audiogenic seizure severity as follows: 0 = no observed seizure activity, 1 = progression to convulsions, 2 = progression to respiratory failure. N = 12 WT, 8 WT;Smad4 cKO, 15 FXS, 16 FXS;Smad4 cKO mice. Statistics by Chi-squared pairwise between groups, using counts in each of the three categories. Chi-squared for 4 groups: p = 0.0045. **h**. Smad4 cKO reduces respiratory failure from audiogenic seizures, same data as **g**. replotted. N = 12 WT, 8 WT;Smad4 cKO, 15 FXS, 16 FXS;Smad4 cKO mice. Pairwise statistics by Fisher’s exact test, using counts. Chi-squared for 4 groups: p = 0.0279.

In the present study, we remove Smad4 from astrocytes to suppress astrocyte BMP signaling *in vivo*, and ask whether a Smad4 conditional KO (cKO) can improve Fmr1 KO phenotypes. We find that Smad4 cKO lessens Fmr1 KO audiogenic seizure severity but does not improve Fmr1 KO visual acuity or ocular dominance plasticity deficits. To gain a detailed unbiased understanding of molecular pathways changed in Fmr1 KO and Smad4 cKO that may underlie these effects, we perform multi-omic astrocyte profiling. We find transcriptional evidence for a hypermetabolic state in Fmr1 KO astrocytes with partial correction in Smad4 cKO. We quantify astrocyte-secreted and membrane proteins *in vivo*, and find in Fmr1 KO a downregulation of vesicular trafficking protein machinery, with partial correction in Smad4 cKO. Functionally, Smad4 cKO rescues an Fmr1 KO decrease in auditory cortex inhibitory synaptic density. Our findings provide new resources for interrogating astrocyte state in Fmr1 KO and astrocyte BMP signaling, deepen the understanding of astrocyte involvement in FXS, and delineate the extent to which astrocyte BMP signaling underlies Fmr1 KO molecular and functional phenotypes.

## Results

### Smad4 cKO suppresses *in vivo* astrocyte BMP signaling

Our hypothesis is that upregulated BMP signaling in astrocytes contributes to alterations in FXS. To suppress BMP signaling specifically in astrocytes *in vivo*, we utilized an astrocyte-specific Smad4 conditional knockout (Smad4 cKO) by crossing Smad4 floxed mice^27^ to the astrocyte-specific Gfap-Cre73.12 line^28,29^ (**Fig. 1B**). Given the presence of visual and auditory manifestations in FXS patients,^1,30^ and the use of visual (acuity,^31^ plasticity^32^) and auditory (audiogenic seizure^33^) phenotypes in Fmr1 KO research, we focused our validation on the visual and auditory pathways. Immunohistochemistry for SMAD4, astrocyte nuclear marker SOX9, and neuron nuclear marker NEUN in P28 auditory and visual cortices showed 68±7% and 83±4% decrease in astrocyte nuclei immunopositive for SMAD4 respectively with no change in neuronal nuclei SMAD4 immunopositivity (**Fig. 1C,D,E, Supplementary Fig. 1B,D**). Moreover, the Smad4 cKO was effective across cortical layers, with each layer of auditory and visual cortex having >64% reduction of SMAD4 positive SOX9+ cells (**Supplementary Fig. 1A,C**), and moderately effective in the inferior and superior colliculi, auditory and visual recipient zones respectively, with 16-26% decrease overall and 37-57% decrease in superficial areas (**Supplementary Fig. 1E,F**). To ensure that SMAD4 was removed from astrocytes during early development, we did further analysis at P14, finding a decrease in SMAD4 in astrocytes by immunostaining in auditory and visual cortices, as well as by other approaches (**Supplementary Fig. 2**).

To validate that the Smad4 cKO effectively suppresses BMP signaling in astrocytes, we performed immunohistochemistry for the known BMP-induced protein ID3.^15^ ID3 expression in SOX9+ astrocyte nuclei in P28 cortex was significantly downregulated in Smad4 cKO (**Supplementary Fig. 1G,H**). Given the known role for BMP signaling in astrocyte cell fate and neurogenesis inhibition,^16,34^ and a recent report of Smad4 involvement in astrocyte proliferation,^35^ we also quantified cortical layer thickness, astrocyte number, and total cell number in P28 cortex. We found no difference in astrocyte or total cell numbers in Smad4 cKO mice relative to littermate controls, and a small decrease in cell density (**Supplementary Fig. 3B,F**) driven by a small increase in cortical thickness (**Supplementary Fig. 3D,H**), indicating no gross alteration in cell fate. Overall, we generated and validated astrocyte Smad4 cKO, a mouse model with specific suppression of BMP signaling in astrocytes.

### Smad4 cKO reduces Fmr1 KO audiogenic seizure severity

With a model validated to suppress BMP signaling in astrocytes, we first asked if suppressing astrocyte BMP signaling can rescue FXS functional phenotypes. We tested for improvement in audiogenic seizures (AGS), a prevalent assay in Fmr1 KO research that models FXS patient hypersensitivity and seizures.^33^ In this assay, P20-23 mice in the four genotypes (1) Fmr1 WT;Smad4 WT (WT), (2) Fmr1 WT;Smad4 cKO (WT;Smad4 cKO), (3) Fmr1 KO;Smad4 WT (FXS), and (4) Fmr1 KO;Smad4 cKO (FXS;Smad4 cKO) were tested for susceptibility to and severity of audiogenic seizures (**Fig. 1B,F**). Seizures were scored on a 0-2 scale of increasing severity: (0) no reaction, (1) convulsions, and (2) respiratory failure. We used male mice because female FXS patients (and mice) are heterozygous and mosaic for the FMR1 gene, leading to less severe and more variable manifestiations.^36^ We found that FXS mice had more severe seizures than WT mice, and that FXS;Smad4 cKO mice had less severe seizures than FXS mice (**Fig. 1G**). Importantly, FXS;Smad4 cKO mice had a significantly lower proportion of mice with score 2 (1 of 16) than FXS mice (8 of 15) (**Fig. 1H**). Smad4 cKO thus reduces Fmr1 KO audiogenic seizure severity.

We next asked if Smad4 deletion in astrocytes could similarly rescue visual Fmr1 KO phenotypes. Visual phenotypes of Fmr1 KO mice include delayed learning in a visual discrimination task that models FXS patient visual discrimination deficits^30^ and increased ocular dominance plasticity (ODP).^32^ We elected to probe Fmr1 KO mouse vision with the optomotor task for visual acuity,^37,38^ and found that at P26, Fmr1 KO mice have decreased visual acuity relative to WT littermates (WT: 0.376±0.002 cyc/deg, Fmr1 KO: 0.349±0.007 cyc/deg) (**Supplementary Fig. 4A**). However, Smad4 cKO did not improve Fmr1 KO decreased visual acuity (FXS: 0.354±0.005 cyc/deg; FXS;Smad4 cKO: 0.349±0.007 cyc/deg) (**Supplementary Fig. 4B**). To test for ocular dominance plasticity, we assayed for binocular zone enlargement during the visual critical period after monocular enucleation (ME) via single molecule fluorescence *in situ* hybridization for immediate early gene *Arc*, as described previously^39^ (**Supplementary Fig. 4C**). Paralleling the known Fmr1 KO ocular dominance hyperplasticity phenotype^32^ and the delayed critical period phenotype across Fmr1 KO sensory systems,^40^ we found that in Fmr1 KO, the size of the activation zone after 4 days visual deprivation was significantly larger than in WT littermate pairs (WT: 1180±55μm; FXS: 1390±51μm) (**Supplementary Fig. 4D,E**). However, Smad4 cKO did not significantly change activation zone width in Fmr1 KO mice (**Supplementary Fig. 4F**), indicating no improvement in Fmr1 KO hyperplastic ocular dominance plasticity. Overall, we validated three functional Fmr1 KO phenotypes, and found that Smad4 cKO in astrocytes does not change Fmr1 KO visual acuity or ocular dominance plasticity, but specifically reduces Fmr1 KO audiogenic seizure severity.

### Fmr1 KO astrocytes have a hypermetabolic transcriptional state *in vivo*

Given that suppressing astrocyte BMP signaling can reduce some FXS functional deficits, we sought to identify how this occurs on a molecular and pathway level. We first used RNA sequencing to unbiasedly identify which specific astrocyte gene programs are altered in Fmr1 KO, and to ask if suppression of astrocyte BMP signaling improves them. We profiled cortical astrocytes at P26-28 in mice of the four genotypes (1) WT, (2) WT;Smad4 cKO, (3) FXS, and (4) FXS;Smad4 cKO (N = 4-6 mice/genotype) (**Fig. 2B**). We used whole cortex in order to obtain enough astrocytes to perform bulk RNA sequencing, and isolated astrocyte nuclei using glyoxal fixation combined with antibody staining for the astrocyte marker SOX9 and fluorescence-activated cell sorting (FACS) purification (**Fig. 2A**, **Supplementary Fig. 5A**).^41,42^ To validate astrocyte specificity of purified nuclei, we averaged the transcript per million (TPM) for a series of known cell type-specific genes,^43^ and found enrichment for astrocytes over other cell types (**Supplementary Fig. 5B**). Inspection of aligned sequencing reads to *Smad4* exon 9 and *Fmr1* exon 5 loci were consistent with Smad4 recombination and Fmr1 KO respectively (**Supplementary Fig. 5C**). As further confirmation for the data in **Fig. 1** showing suppression of BMP signaling, we saw a decrease in *Id3* expression in Smad4 cKO (p-adj = 3.1e-20, log2 fold change [L2FC] = -2.28; **Supplementary Data Table 2**).

**Figure 2.**
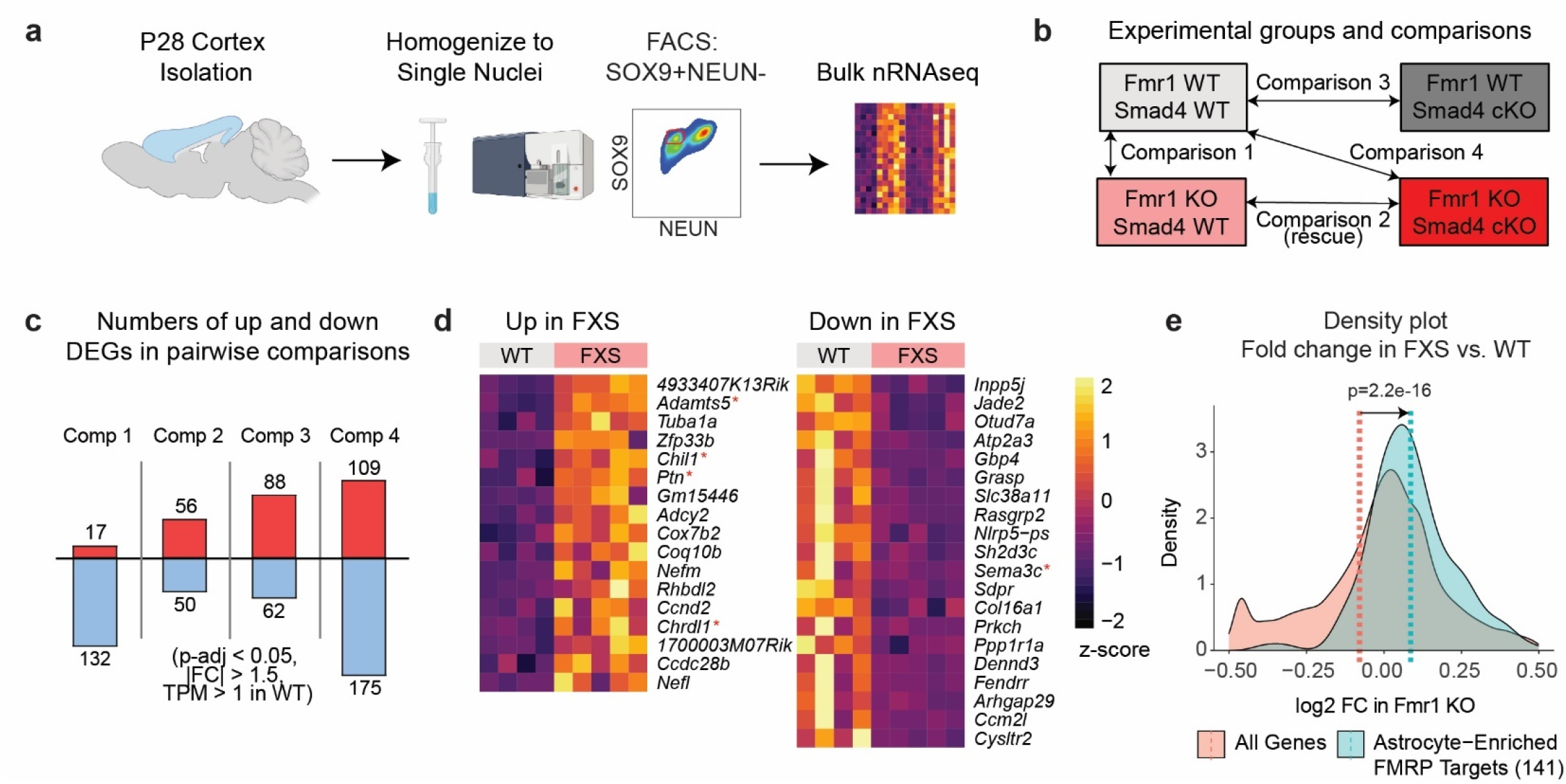
Nuclear transcriptomics identifies Fmr1 KO and Smad4 cKO astrocyte *in vivo* transcriptomic dysregulation. **a.** Protocol for glyoxal-fixed astrocyte nuclei transcriptomics via FACS sorting for SOX9+NEUN-nuclei followed by bulk nRNAseq. **b**. Experimental groups and pairwise comparisons. **c**. Numbers of up- and down-DEGs for the transcriptomic experiment. N = 4 WT, 6 WT;Smad4 cKO, 5 FXS, 4 FXS;Smad4 cKO mice. Cutoffs p-adj < 0.05, |FC| > 1.5, TPM in WT > 1 calculated with DESeq2 with Benjamini-Hochberg (BH) correction. See also Supplementary Data Tables 1,2. **d**. Top significantly up- and downregulated genes in Fmr1 KO (Comparison 1). Genes filtered by |FC| > 1.5 and TPM in WT > 1, sorted by p-adj. Red asterisks indicate genes encoding known astrocyte-secreted proteins. **e**. Density plot of log2 fold change in Fmr1 KO (Comparison 1) for two gene sets: (1) all genes detected in nRNAseq, (2) FMRP targets (842 genes, Darnell et al. 2011^52^), filtered to those enriched at an mRNA level in astrocytes (141 genes, Zhang et al. 2014^53^). The average log2 fold changes for these gene sets are denoted with dashed lines. The two distributions are significantly different, statistics by z-test.

We compared FXS to WT (Comparison 1) to determine genes dysregulated by Fmr1 KO in astrocytes, finding 149 differentially expressed genes (DEGs) (**Fig. 2C**, **Supplementary Data Table 2**; threshold p-adj < 0.05, |fold change (FC)| > 1.5, TPM in WT > 1). Interestingly, genes encoding for astrocyte-secreted proteins were among the top upregulated (*Chil1*, *Ptn*, *Chrdl1*) and downregulated (*Sema3c*) when ranked by Benjamini-Hochberg (B-H) adjusted p-value (**Fig. 2D**).^12,39,44,45^ Gene set enrichment analysis (GSEA) revealed top Fmr1 KO upregulated gene sets to be “oxidative phosphorylation,” “cholesterol homeostasis,” and “mTorc1 signaling,” while top downregulated gene sets were related to “interferon response” (Comparison 1, **Fig. 3E**, **Supplementary Data Table 3**). Several groups have reported metabolic hyperactivity in bulk brain tissue and other cell types in Fmr1 KO^1,46–48^ as well as cholesterol dysregulation in Fmr1 KO astrocytes;^49–51^ our data also finds this hypermetabolic transcriptional signature in Fmr1 KO astrocytes *in vivo*.

**Figure 3.**
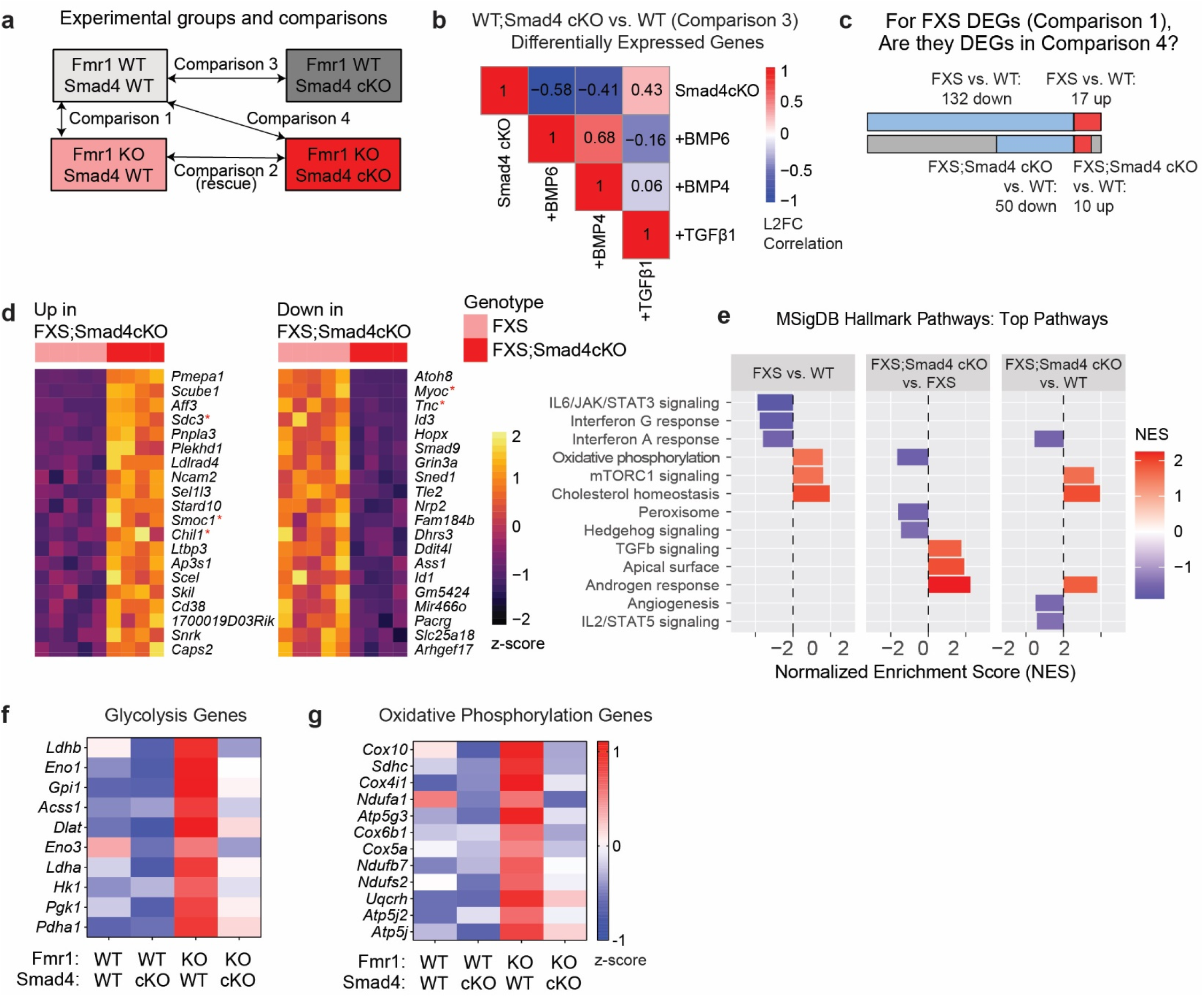
Smad4 cKO moderates Fmr1 KO astrocyte transcriptomic activation of metabolic pathways. **a**. Pairwise comparisons and groups for the RNAseq experiment. **b**. Correlation matrix for L2FCs of the 150 *in vivo* RNAseq Smad4 cKO DEGs, between *in vivo* RNAseq of Smad4 cKO, WT astrocytes incubated with BMP6,^12^ and cultured astrocytes incubated with BMP4 or TGFβ1.^54^ **c**. For the 149 FXS DEGs (Comparison 1), the majority are no longer significantly changed in Comparison 4 (FXS;Smad4 cKO vs. WT). See Supplementary Data Table 2 for gene list. **d**. Top significantly up- and downregulated genes in FXS;Smad4 cKO compared to FXS (Comparison 2). Genes filtered by |FC| > 1.5 and TPM in WT > 1, sorted by p-adj. Red asterisks indicate genes encoding known astrocyte-secreted proteins. **e**. Top 3 significantly up- and down-regulated pathways by normalized enrichment score (NES), for each of the comparisons 1, 2, and 4; via gene set enrichment analysis on the MSigDB Hallmark Gene Sets with all genes TPM > 1 ranked by –log10(p-value) * sign(L2FC). See also Supplementary Data Table 3. **f-g**. Z-scores of genes in **f**. “glycolysis/gluconeogenesis” (KEGG M11521) and **g**. “oxidative phosphorylation” (MSigDB MM3893) gene sets, averaged within each genotype; displaying genes up in Fmr1 KO and lowered by Smad4 cKO. See Supplementary Fig. 5d,e for full gene sets.

Additionally, we compared these Fmr1 KO DEGs to a list of 842 genes known to be bound to and thus likely regulated by fragile X messenger ribonucleoprotein (FMRP), the protein product of *Fmr1*.^52^ While only 4 DEGs (*Kcnt1, Plxnd1, Rn45s, Shank3*) were known FMRP targets (p = 0.76), overall, the set of FMRP targets filtered to those enriched in astrocytes at an mRNA level^53^ (141 genes) had an average log2 fold change (FXS vs. WT) greater than the set of all genes (p = 2.2e-16, **Fig. 2E**), implying FMRP directly regulates gene expression in astrocytes. We also compared this *in vivo* nuclear RNA transcriptomic dataset to an *in vivo* P40 Fmr1 KO astrocyte translating RNAseq dataset,^11^ and a cultured P7 Fmr1 KO astrocyte transcriptome,^12^ and surprisingly found low overlap and minimal correlation in log2 fold change of DEGs (**Supplementary Fig. 6A,C,D,E**), highlighting the roles that cellular context and timepoint may play in identifying transcriptomic alterations in a disease model.

### Smad4 cKO moderates Fmr1 KO astrocyte transcriptomic activation of metabolic pathways

Our primary goal was to determine if Smad4 cKO can rescue Fmr1 KO astrocyte dysregulation, and elucidate molecular pathways underlying any functional improvement. We thus performed the additional pairwise comparisons: WT;Smad4 cKO to WT (Comparison 3), FXS;Smad4 cKO to FXS (Comparison 2), and FXS;Smad4 cKO to WT (Comparison 4) (**Fig. 3A**). First, examining the effect of Smad4 cKO in WT (Comparison 3), we found 150 DEGs (**Supplementary Data Table 2**). Interestingly, log2 fold changes (L2FC) of Smad4 cKO DEGs correlated negatively to L2FCs of the same genes in astrocytes cultured with BMP4 (r = -0.41) or BMP6 (r = -0.60) and not TGFβ1 (r = +0.39) (**Fig. 3B**),^54^ suggesting preferential suppression of the astrocyte BMP and not TGFβ transcriptional signature in Smad4 cKO. Moreover, at the level of individual DEGs, 44% of downregulated DEGs in Smad4 cKO *in vivo* were upregulated DEGs in astrocytes cultured with BMP6 (**Supplementary Fig. 6B**), showing similarity in astrocyte transcriptomic response to BMP signaling *in vitro* and *in vivo*.

Comparing FXS;Smad4 cKO to WT (Comparison 4), we found 284 DEGs (**Fig. 2C, Supplementary Data Table 2**). Notably, the majority (89 of 149) of DEGs found when comparing FXS to WT (Comparison 1) were no longer significantly changed when comparing FXS;Smad4 cKO to WT (Comparison 4) (**Fig. 3C**, **Supplementary Data Table 2**). These 89 genes thus represent FXS astrocyte transcriptional dysregulations moderated by Smad4 cKO. To directly assess the impact of Smad4 cKO in the FXS background, we also compared FXS;Smad4 cKO to FXS (Comparison 2). We found 106 DEGs, and again, genes encoding astrocyte-secreted proteins were among top upregulated (*Sdc3, Smoc1*) and downregulated (*Myoc*, *Tnc*) genes (**Fig. 3D, Supplementary Data Table 2**).

At the pathway level, strikingly, GSEA showed “oxidative phosphorylation” is a top suppressed pathway comparing FXS;Smad4 cKO to FXS (Comparison 2), and is not significantly changed comparing FXS;Smad4 cKO to WT (Comparison 4), demonstrating Smad4 cKO has rescued this FXS alteration (**Fig. 3E**). This is in contrast to the “mTORC1 signaling” and “cholesterol homeostasis” pathways, which are upregulated in Fmr1 KO and not impacted by Smad4 cKO, and the “interferon A response” pathway, which is downregulated in Fmr1 KO and not impacted by Smad4 cKO. GSEA analysis on a broader set of gene sets showed “glycolysis” also followed the same trend as “oxidative phosphorylation,” with upregulation of this pathway in FXS moderated by Smad4 cKO (**Fig. 3F,G**, **Supplementary Fig. 5D,E**, **Supplementary Data Table 3,4**). The effect we see in Smad4 cKO is consistent with reports in other cell types showing positive regulation of glycolysis and oxidative phosphorylation by BMP signaling.^55^ In summary, our RNAseq data demonstrate dysregulation of the astrocyte transcriptome in Fmr1 KO, and we determine a number of genes and pathways partially rescued by Smad4 cKO including the hypermetabolic phenotype present in FXS astrocytes.

### Interrogating *in vivo* astrocyte endoplasmic reticulum-passaged proteins

Astrocyte-secreted proteins are known to contribute to *in vitro* Fmr1 KO phenotypes.^12^ Intriguingly, from the above RNAseq, genes encoding known astrocyte-secreted proteins^12,39,45,56,57^ were among the top Fmr1 KO upregulated (*Adamts5*, *Chil1, Ptn*, *Chrdl1*) and downregulated (*Sema3c*) genes (**Fig. 2D**), as well as top Smad4 cKO upregulated (*Sdc3*, *Sparc, C4b*) and downregulated (*Myoc*, *Tnc*, *Serpinf1*) genes (**Fig. 3D**, **Supplementary Data Table 2**). However, correlation between astrocyte transcriptional changes and protein secretion changes is limited,^12^ and we thus sought to quantify *in vivo* changes in astrocyte secretion at the protein level.

To profile the full complement of astrocyte-secreted proteins *in vivo*, we took a proximity labeling proteomic approach with the enzyme TurboID,^58^ which labels nearby proteins with biotin. Expressing TurboID in a cell type- and compartment-specific manner enables *in vivo* biotin tagging and subsequent streptavidin enrichment of proteins within a specific cellular compartment. We subcloned and tested three constructs based on subcellular TurboID localizations known to target secreted proteins: endoplasmic reticulum membrane (ERM) facing cytosol (via cytochrome P450 (C1(1-29)) targeting sequence),^58^ ERM facing ER lumen (via fusion protein with Sec61b),^59^ and ER lumen (via KDEL ER retention signal) (**Supplementary Fig. 7A**).^59,60^ While all three constructs were functional *in vitro* (**Supplementary Fig. 7B**), pilot mass spectrometry on astrocyte conditioned media and lysates suggested highest labeling of secreted proteins by the ER lumen construct (ER-TurboID). To test ER-TurboID *in vivo*, we packaged the construct into AAV-PHP.eB (an adeno-associated virus with serotype allowing crossing of the blood-brain barrier to transduce the CNS),^61^ injected virus retroorbitally at P12, labeled with exogenous biotin by subcutaneous injection from P21 to P26, and collected mice at P27 (**Fig. 4A**). ER-TurboID was highly expressed by P20 (**Supplementary Fig. 8A**) and incorporated biotin efficiently and specifically in astrocytes at P27 (**Fig. 4B,C**, **Supplementary Fig. 8D**). Thus we adapted and optimized a protocol to label astrocyte-secreted proteins *in vivo*.

**Figure 4.**
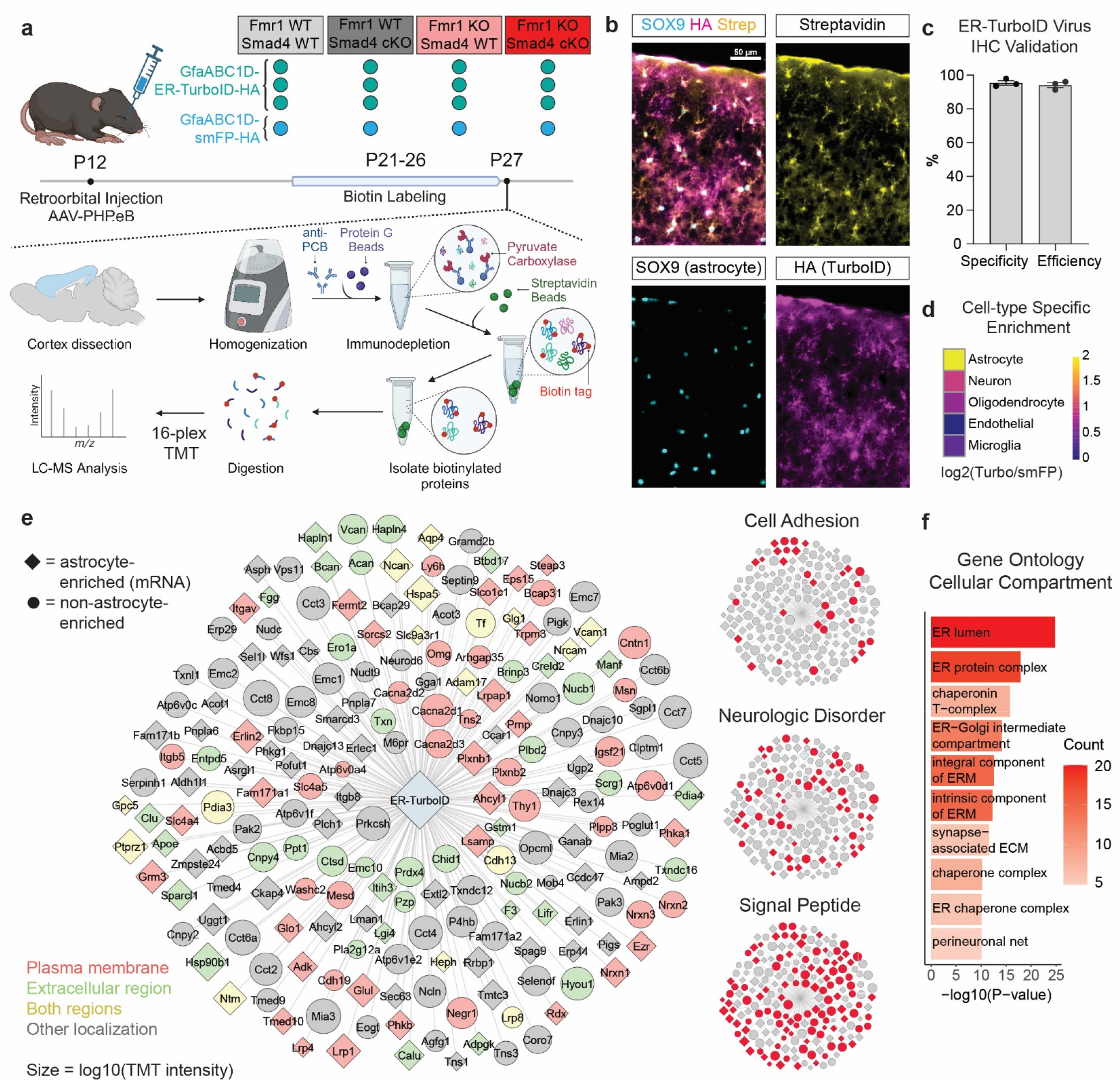
Development of ER-targeted TurboID for *in vivo* astrocyte secreted and membrane proteomics. **a**. Protocol for ER-TurboID proteomics. **b**. ER-TurboID biotinylates proteins in SOX9+ astrocytes. Immunohistochemistry imaged in visual cortex, for a mouse subjected to the protocol in **a**. with fixation instead of dissection for proteomics. HA tag on ER-TurboID construct colocalizes with streptavidin (biotinylation) in cells with SOX9+ nuclei. **c**. Quantification of **b**. Specificity quantified as (# SOX9+HA+ cells)/(# HA+ cells), efficiency quantified as (# SOX9+HA+ cells)/(# SOX9+ cells) for N = 3 mice. **d**. ER-TurboID enriches for known astrocyte proteins. Plotting log2 of average TurboID to average control TMT signal, averaged across known cell type-specific proteins (see Supplementary Fig. 8e for all proteins). **e**. Most highly enriched proteins in the ER-TurboID proteome (log2[Turbo/Control] > 2.38, 200 proteins). Node tile size represents log10 of average TMT signal for the protein across all experimental samples. Tile shape represents cell type: diamond = astrocyte (mRNA level higher in astrocytes compared to the average expression in other cell types^53^), circle = other cell type or unknown. Tile color represents cellular compartment gene ontology annotation: green = extracellular region, red = plasma membrane, yellow = both extracellular region and plasma membrane, grey = other. Right, highlighting proteins in the functional category of cell adhesion (GO: 7155), with causative mutation linked to neurologic disorder, and with predicted signal peptide. **f**. Gene ontology overrepresentation analysis based on the 200 proteins in **e**., listing the top 10 cellular compartment categories by p-value.

To identify astrocyte-secreted proteins changed in Fmr1 KO and Smad4 cKO, three mice each of the four genotypes (1) WT, (2) WT;Smad4 cKO, (3) FXS, and (4) FXS;Smad4 cKO were injected with ER-TurboID virus (**Fig. 4A**). One mouse of each genotype was injected with virus expressing control protein spaghetti monster fluorescent protein (smFP-HA, **Fig. 4A**, **Supplementary Fig. 8B,C**). All mice were given exogenous biotin, cortices were lysed, depleted for endogenously biotinylated pyruvate carboxylase,^62^ and biotinylated proteins were purified by streptavidin pulldown, pooled for 16-plex tandem mass tagging (TMT) to enable quantification using isotopic labeling, and analyzed by high-resolution liquid chromatography-mass spectrometry (LC-MS) (**Fig. 4A**). In total MS identified 22609 peptides corresponding to 2138 distinct proteins (**Supplementary Data Table 6**). To validate enrichment of astrocyte labeling, we took detected proteins from a list of cell type-specific markers,^43^ then calculated for each protein the ratio of (1) the average TMT signal across the 12 ER-TurboID samples, compared to (2) the average TMT signal across the 4 control samples. Astrocyte-specific markers showed highest enrichment compared to other cell type markers (log2 Turbo/Control ratio = 2.00), validating this approach (**Fig. 4D**, **Supplementary Fig. 8E**).

To better understand the proteins labeled by ER-TurboID, we took the 200 proteins with highest enrichment (corresponding to log2[Turbo/Control] > 2.38) to characterize the astrocyte ER-TurboID proteome (**Fig. 4E**). This list includes known astrocyte-secreted proteins (e.g. APOE, GPC5, CLU, SPARCL1) as well as known astrocyte membrane proteins (e.g. ITGB5, GRM3, AQP4) and many perineuronal net (PNN) components (e.g. ACAN, BCAN, NCAN, VCAN, HAPLN1, HAPLN4). Cross-referencing a cell type-specific gene expression database showed a substantial number of proteins have known mRNA-level enrichment in astrocytes relative to other CNS cell types (95/200, 48%) (**Fig. 4E**). Bioinformatics analysis showed that this proteome contains proteins localizing to the extracellular region (55 of 200, 28%), plasma membrane (67 of 200, 34%), or both (16/200, 8%); proteins predicted to have a signal peptide (111/200, 56%); and proteins encoded by genes implicated in neurologic disorders (57/200, 29%) (**Fig. 4E**). Moreover, gene ontology analysis for cellular compartment showed these 200 proteins are significantly enriched for the endoplasmic reticulum, membrane, and extracellular compartments (**Fig. 4F**). In summary, we adapted and validated a proximity labeling approach to quantify astrocyte-secreted and membrane proteins *in vivo*.

### Fmr1 KO astrocytes downregulate extracellular matrix proteins and secretory machinery

To determine proteins dysregulated in Fmr1 KO astrocytes, we compared FXS to WT (Comparison 1), finding 95 differentially expressed proteins (DEPs) (**Fig. 5A**, **Supplementary Data Table 7**; threshold p < 0.05, |fold change (FC)| > 1.25), 97% of which were enriched in TurboID over control. Among the Fmr1 KO DEPs (**Fig. 5B,D**), we found downregulation of many perineuronal net components (ACAN, BCAN, NCAN, HAPLN1, HAPLN4, TNR), in concordance with Fmr1 KO literature documenting immature perineuronal nets in brain regions including auditory cortex.^63–65^ Additionally, many cell membrane proteins with known genetic association to autism spectrum disorder were downregulated in Fmr1 KO (e.g. NRXN2,^66^ NRXN3,^67^ PLXNB1,^68^ LRP1,^68^ NRCAM^69^). Moreover, Fmr1 KO downregulated DEPs overlapped significantly with known FMRP targets^52^ (17/89 = 19% overlap, p = 9.6e-9) (**Fig. 5C**), demonstrating that FMRP regulates protein expression in astrocytes. Cellular compartment analysis of the Fmr1 KO DEPs highlights that many detected DEPs localize to the ER, vesicles, membrane, and extracellular compartments, and have signal peptides (**Fig. 5E,F**), as expected for this ER-TurboID proteomic method. At the pathway level, GSEA in the Fmr1 KO vs WT comparison (Comparison 1) showed downregulation of gene sets “protein secretion” and “apical junction” and upregulation of “mitotic spindle,” among other changes (**Fig. 6D**, **Supplementary Data Table 8**).

**Figure 5.**
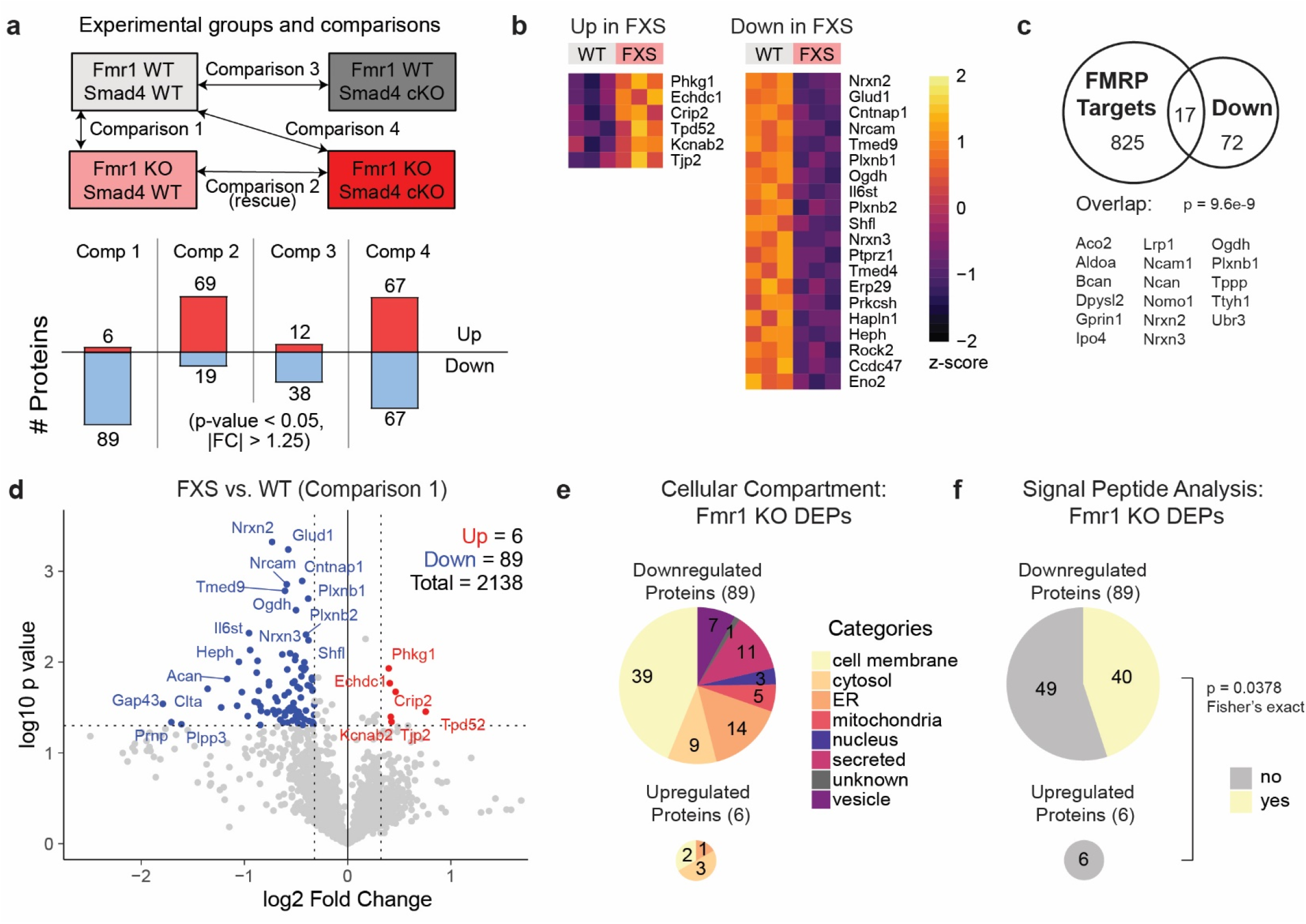
ER-TurboID identifies dysregulated Fmr1 KO astrocyte secreted and membrane proteins. **a**. Above, groups and comparisons for the proteomic experiment. N = 3 WT, 3 WT;Smad4 cKO, 3 FXS, 3 FXS;Smad4 cKO mice. Proteomics processed with 16-plex TMT, including one mouse per genotype injected with virus with plasmid expressing control protein. Below, number of up- and downregulated proteins for each of the 4 comparisons, cutoffs p < 0.05, |FC| > 1.25. See also Supplementary Data Tables 6,7. **b**. Top up- and downregulated proteins in Fmr1 KO (Comparison 1). Proteins sorted by p-value. **c**. Overlap of known FMRP targets^52^ with downregulated proteins in Fmr1 KO (Comparison 1). Statistics by Fisher’s exact test with genome size 24,252 protein-coding *M. Musculus* genes. **d**. Volcano plot for the Fmr1 KO vs WT comparison showing significantly up- and downregulated proteins. Dashed black lines at p = 0.05 and |FC| = 1.25. **e**. Cellular compartment and **f**. signal peptide analysis of DEPs from **d**., based on UniProt annotations.

**Figure 6.**
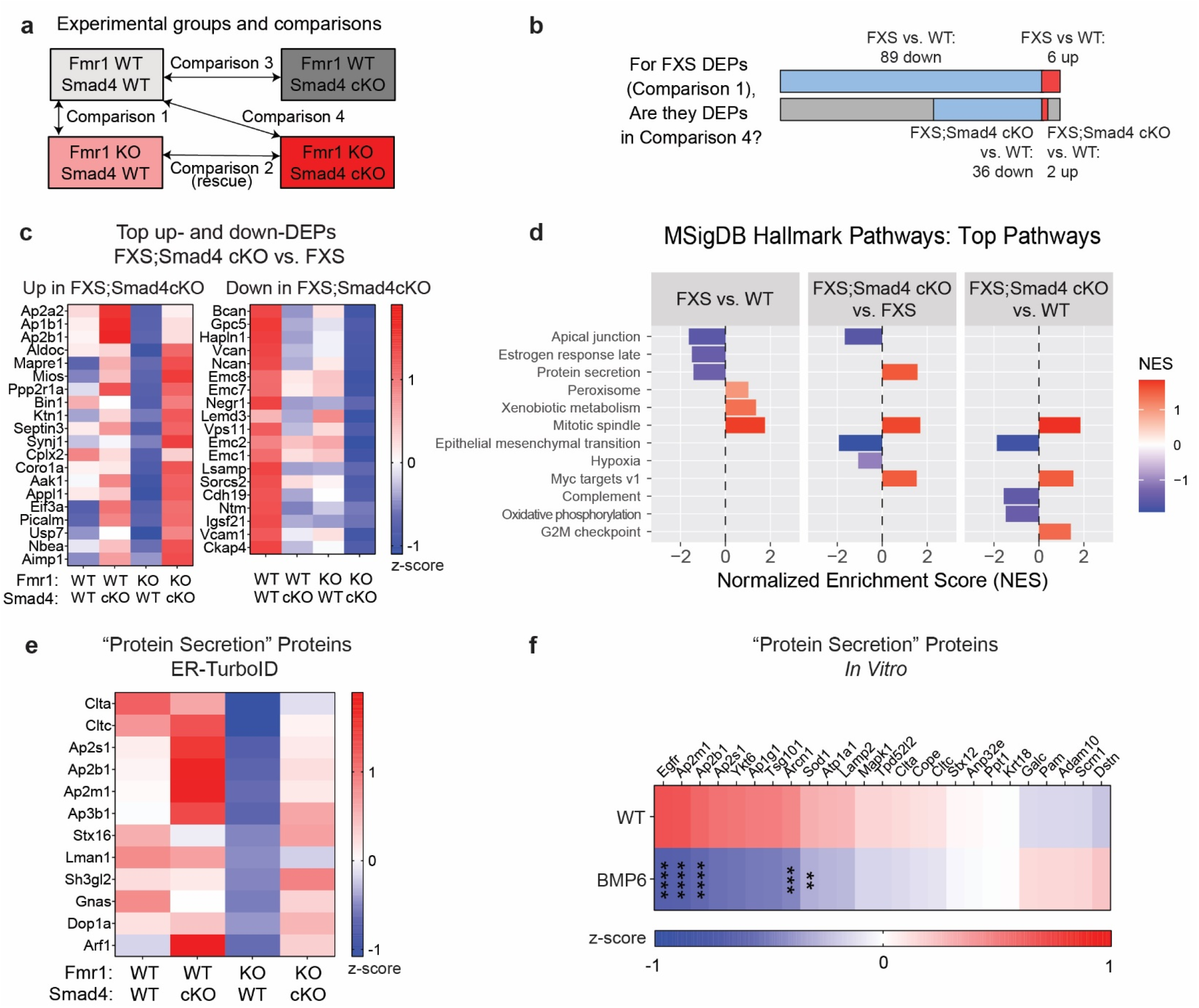
Smad4 cKO restores Fmr1 KO astrocyte deficit in secretory machinery. **a**. Groups and comparisons for the proteomic experiment. **b**. For the 95 FXS DEPs (Comparison 1), the majority are no longer significantly changed in Comparison 4 (FXS;Smad4 cKO vs. WT). See Supplementary Data Table 7 for protein list. **c**. Average z-scores by genotype of the top significantly up- and downregulated proteins in FXS;Smad4 cKO compared to FXS (Comparison 2). Proteins sorted by p-value. **d**. Top 3 up- and down-regulated pathways by normalized enrichment score (NES), for each of the comparisons 1, 2, and 4; via gene set enrichment analysis on the MSigDB Hallmark Gene Sets with all proteins Turbo/Control > 1.75 ranked by –log10(p-value) * sign(L2FC). See also Supplementary Data Table 8. **e**. Z-scores of proteins in the “protein secretion” gene set (MSigDB MM3876), averaged within each genotype; displaying proteins down in Fmr1 KO and increased by Smad4 cKO. See Supplementary Fig. 12 for full set. **f**. Protein secretion machinery is downregulated in the secretome of P7 WT astrocytes cultured with BMP6, data re-analyzed from Caldwell et al., 2022.^12^ Z-scores of all detected proteins in the “protein secretion” gene set (MSigDB MM3876), averaged within each genotype. Statistics as in Caldwell et al., 2022.^12^ ** denotes p < 0.01, *** denotes p < 0.001, **** denotes p < 0.0001.

We asked to what extent the changes we found in gene expression (RNAseq) were recapitulated at the protein level (ER-TurboID). We found no overlap between DEGs and DEPs, and Fmr1 KO vs WT fold changes for TPM (RNA) and TMT signal (protein) showed low correlation (**Supplementary Fig. 9A,B,C**). At the level of absolute abundance, there was a low, slightly positive, correlation (**Supplementary Fig. 9D,E,F**). This low degree of overlap between RNA and protein findings may be due to the different cellular compartments assayed (nuclear RNA vs. ER-transited proteins)^70^ as well as the uncoupling of transcription and translation in the absence of translational regulator FMRP,^71^ but also illustrates the importance of transcriptomics and proteomics as independent readouts of cellular state and function. Finally, we compared this *in vivo* proteomic dataset to an *in vivo* P40 Fmr1 KO astrocyte whole-cell proteomics dataset^11^ and a cultured P7 Fmr1 KO astrocyte secreted protein dataset^12^ and surprisingly found low overlap between the DEPs, and minimal correlations in log2 fold change of DEPs (**Supplementary Fig. 10A,C,D,E**), highlighting the roles that cellular context, subcellular compartment, and timepoint may play in proteomic variability.

### Smad4 cKO restores Fmr1 KO astrocyte deficit in secretory machinery

Our primary goal was to determine if Smad4 cKO can rescue Fmr1 KO astrocyte dysregulation, and elucidate pathways underlying functional rescues. We thus asked if Smad4 cKO could rescue Fmr1 KO astrocyte proteomic dysregulation by performing pairwise comparisons across the four genotypes (1) WT, (2) WT;Smad4 cKO, (3) FXS, and (4) FXS;Smad4 cKO. First, to validate our proteomics results, we performed western blotting on cortical lysates from all four genotype groups for brevican (BCAN), a protein downregulated in the ER-TurboID data in FXS and Smad4 cKO (**Supplementary Data Table 6,7**) that is known to be astrocyte-enriched,^53^ BMP-induced,^54,72^ and a component of perineuronal nets. We found the same changes in BCAN expression across genotypes in western blot and ER-TurboID proteomics: a decrease in both Smad4 cKO and Fmr1 KO (**Supplementary Fig. 11A,B,C**).

We then examined differentially expressed proteins and pathways in the pairwise comparisons FXS;Smad4 cKO to FXS (Comparison 2) and FXS;Smad4 cKO to WT (Comparison 4) (**Fig. 6A**). Comparing FXS;Smad4 cKO to WT (Comparison 4), we observed 134 DEPs (**Supplementary Data Table 7**). Notably, the majority (57 of 95) of DEPs found when comparing FXS to WT (Comparison 1) were no longer significantly changed when comparing the FXS;Smad4 cKO to WT (Comparison 4) (**Fig. 6B**, **Supplementary Data Table 7**). These 57 proteins thus represent FXS proteomic dysregulations moderated by Smad4 cKO.

To directly assess the impact of Smad4 cKO in the FXS background, we compared FXS;Smad4 cKO to FXS (Comparison 2). We found 88 DEPs, and saw perineuronal net components as well as astrocyte-secreted Glypican 5 as top downregulated proteins (**Fig. 6C**). In this comparison, top upregulated DEPs in FXS;Smad4 cKO from FXS included both proteins restored to near WT levels (e.g. AP2A2, ALDOC, BIN1) and FXS-independent effects (e.g. MAPRE1, EIF3A) (**Fig. 6C**). In contrast, top downregulated DEPs in FXS;Smad4 cKO augmented FXS changes (**Fig. 6C**).

At the pathway level, GSEA showed “protein secretion” was a top upregulated pathway comparing FXS;Smad4 cKO to FXS (Comparison 2), and not significantly changed when comparing FXS;Smad4 cKO to WT (Comparison 4) (**Fig. 6D**). This reflects a rescue of the FXS downregulation of the “protein secretion” pathway, and was driven by several proteins in secretory/vesicular trafficking/endocytosis pathway machinery downregulated in Fmr1 KO (e.g. CLTA, CLTC, AP2S1, AP2B1, AP2M1) and restored to near WT levels by Smad4 cKO (**Fig. 6E, Supplementary Fig. 12**). Additionally, vesicular trafficking proteins were decreased in conditioned media of astrocytes cultured with BMP6 (**Fig. 6F**),^12^ strengthening the negative association between astrocyte BMP signaling and secretory machinery. However, Fmr1 KO upregulation of the mitotic spindle protein set and downregulation of the apical junction protein set were not improved by Smad4 cKO. In summary, our *in vivo* astrocyte proteomic dataset identified a downregulation of secreted, membrane, and perineuronal net proteins in Fmr1 KO, with an Fmr1 KO deficit in vesicular trafficking machinery improved by Smad4 cKO.

### Smad4 cKO rescues Fmr1 KO decrease in auditory cortex inhibitory synapse density

With a deeper understanding of the molecular pathways altered in Fmr1 KO astrocytes, as well as those improved by Smad4 cKO, we next sought to leverage our multi-omic datasets to further investigate the mechanism of Smad4 cKO reduction of audiogenic seizure severity (**Fig. 1G**). Astrocytes can regulate synapse formation, function, and elimination through their secreted proteins.^73^ Given that the Fmr1 KO deficit we observe in “vesicular trafficking/protein secretion” (**Fig. 6D,E**) is improved by suppressing astrocyte BMP signaling with Smad4 cKO, we hypothesized Smad4 cKO might improve Fmr1 KO pathology at the level of synapses. Moreover, our data shows many changes in synapse-related genes and proteins including in the astrocyte membrane protein hepaCAM, which can induce inhibitory synaptogenesis^74^ and is upregulated in our astrocyte proteomic dataset by Smad4 cKO in both WT and FXS (**Supplementary Fig. 11D**).

To quantify synaptic changes, we performed immunohistochemistry to label both excitatory and inhibitory synapses at P28. Given our finding that Smad4 cKO can reduce Fmr1 KO audiogenic seizure severity (**Fig. 1G**), we focused on auditory cortex and analyzed synapses in L2/3. Excitatory synaptic density, assayed via colocalization of presynaptic marker Vglut1 with postsynaptic marker PSD95, showed no significant changes across genotypes (**Fig. 7A,B**). For inhibitory synaptic density, in Fmr1 KO we found a decrease in colocalization of presynaptic Vgat with postsynaptic Gephyrin (59±6.3% of WT; **Fig. 7C,D**) driven by a decrease in Gephyrin (65±17% of WT) and a trend towards decreased Vgat (83±6.0% of WT). This is consistent with Fmr1 KO literature, in which GABAergic system dysfunction including decreases in inhibitory synaptic density in brain regions^75^ such as auditory cortex^76^ is thought to underlie Fmr1 KO circuit deficits.^77,78^ Strikingly, in Fmr1 KO with Smad4 cKO, average Vgat, Gephyrin, and colocalized Vgat-Gephyrin density were restored to WT levels (Vgat: 100±9.7% of WT; Gephyrin 97±42% of WT; colocalized Vgat-Gephyrin 101±18% of WT; **Fig. 7D**). In conclusion, we find that Smad4 cKO rescues an Fmr1 KO deficit in inhibitory synaptic density in auditory cortex. Reduced inhibition can underlie seizures; as Smad4 cKO can both restore Fmr1 KO inhibitory deficits and reduce Fmr1 KO audiogenic seizures, restoration of inhibition represents a mechanistic bridge from Smad4 cKO molecular changes to improvements in FXS functional phenotypes.

**Figure 7.**
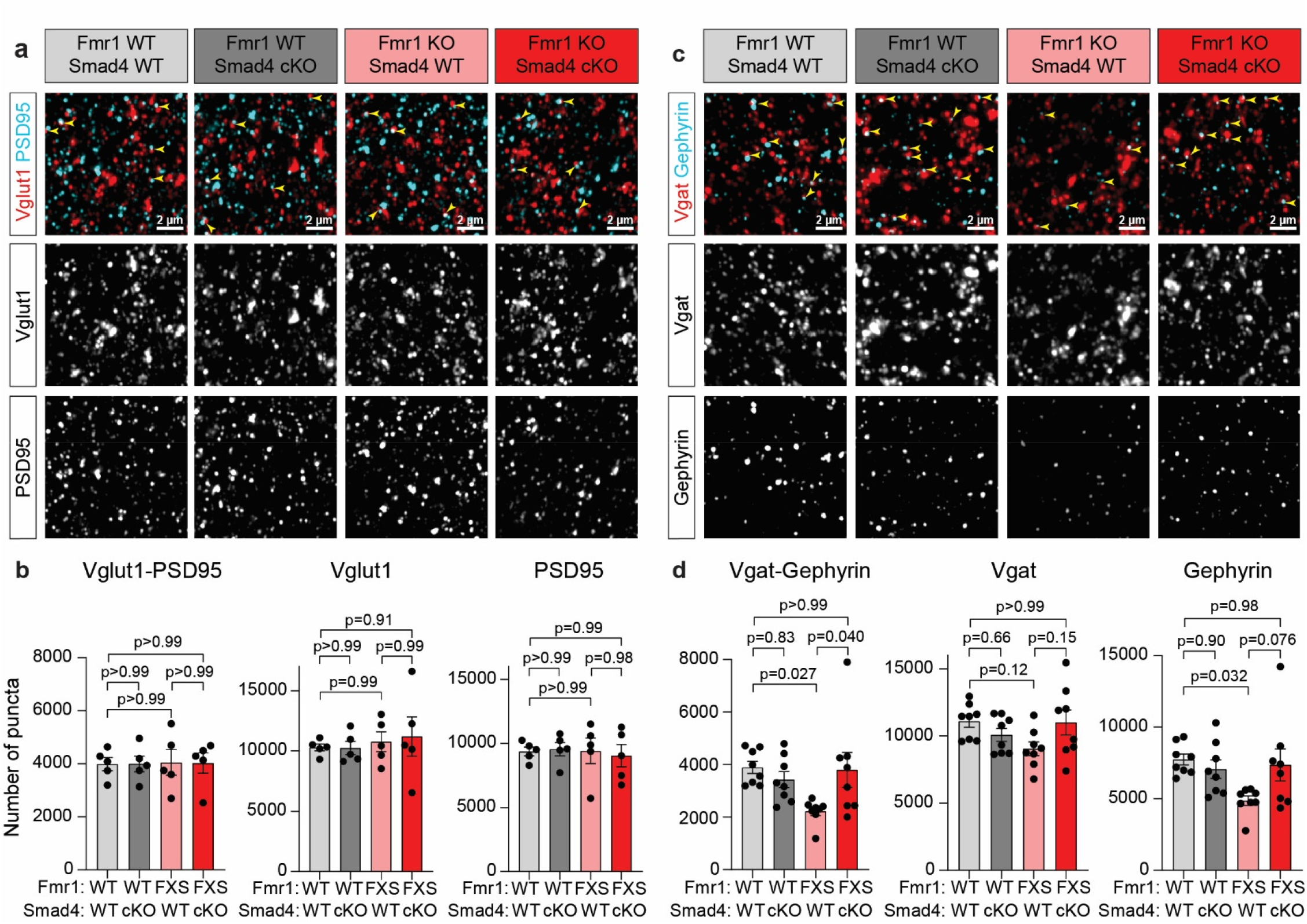
Smad4 cKO rescues Fmr1 KO auditory cortex inhibitory synapse density deficit. **a**. Example excitatory synaptic immunohistochemistry images from L2/3 auditory cortex at P28. Example colocalized Vglut1-PSD95 puncta indicated by yellow arrowheads. **b**. Quantification of **a**., N = 5 mice per genotype. Statistics by 2-way ANOVA with Tukey’s test for multiple comparisons. **c**. Example inhibitory synaptic immunohistochemistry images from L2/3 auditory cortex at P28. Example colocalized Vgat-Gephyrin puncta indicated by yellow arrowheads. **d**. Quantification of **c**., N = 8 mice per genotype. Statistics by 2-way ANOVA with Tukey’s test for multiple comparisons.

## Discussion

In this study we demonstrate that suppressing astrocyte BMP signaling with Smad4 cKO can improve Fmr1 KO phenotypes at multiple levels from molecular to behavioral. This includes correcting molecular signatures at the pathway level such as attenuating astrocyte oxidative phosphorylation upregulation at the transcriptional level and astrocyte secretory machinery downregulation at the proteomic level, as well as improving functional deficits in audiogenic seizure susceptibility and decreased inhibitory synaptic density. We developed a technique for quantification of astrocyte secreted and membrane proteins *in vivo*, and applied this, along with astrocyte-specific transcriptomics, to generate resources to ask how astrocyte state is altered *in vivo* in Fmr1 KO and Smad4 cKO.

Genomic datasets have begun to detail molecular changes caused by loss of FMRP, across the brain and in astrocytes.^11,79^ RNAseq in neurons and bulk tissue and single-cell RNAseq across cortical cell types in Fmr1 KO have shown upregulated translation, mitochondrial processes, and mTOR signaling and downregulated vesicle transport.^80–82^ Single-cell RNAseq in Fmr1 KO cortex also identified dysregulated astrocyte-neuron communication and astrocyte synapse-related genes,^80^ and astrocyte-focused RNAseq has identified hundreds of dysregulated genes *in vitro* and *in vivo* in Fmr1 KO.^11,12^ Our cortical astrocyte transcriptomic profiling showed upregulation of multiple metabolism-related gene sets in Fmr1 KO astrocytes. Astrocytes are highly metabolically active and are known to support neuronal energy balance;^83,84^ conversely, astrocyte metabolic dysfunction is increasingly associated with neurodevelopmental disorders,^47^ with astrocyte cholesterol dysregulation^49,50^ and overproduction of reactive oxidative species (ROS)^85^ tied to FXS. We find that Smad4 cKO moderated Fmr1 KO transcriptional upregulation of oxidative phosphorylation and glycolysis at the pathway level. Our hypothesis that this may underlie rescue of Fmr1 KO phenotypes such as audiogenic seizure susceptibility correlates with other reports that normalizing metabolic dysregulation (RSK signaling,^86^ mitochondrial health,^48,87^ GSK3α ^88^) can normalize Fmr1 KO deficits. Alternatively, the Smad4 cKO rescue of astrocyte metabolic dysregulation in Fmr1 KO could be downstream of improved neuronal hyperexcitability, suggested by the reduced audiogenic seizures and recovered inhibitory synapse levels. This model would explain why a previous *in vitro* transcriptomic study of Fmr1 KO astrocytes cultured in isolation did not show a hypermetabolic signature,^12^ and emphasizes the importance of the cellular environment for phenotypes such as metabolism shaped by intercellular interaction. Indeed astrocytes are known to adapt metabolic flux in response to neuronal activity, both in physiological and pathological contexts.^84^ Further functional studies of *in vivo* astrocyte metabolism in FXS, including with tools beyond the transcriptomics and proteomics employed here, will be crucial to determine if the metabolic effect we observe is primary or secondary in FXS pathophysiology.

Proteomics studies in Fmr1 KO have shown dysregulation in proteins relating to translation machinery, RNA-binding, synaptic plasticity, and mitochondrial function in bulk tissue and in neurons, as well as upregulated complement and coagulation pathways both in bulk tissue and in astrocytes.^12,71,89^ However, until recently there has been an absence of tools to quantify astrocyte-secreted proteins *in vivo*. We developed a new approach to profile astrocyte ER-passaged proteins and quantify secreted and membrane proteins across different conditions during a specified timeframe. We identified that many FMRP targets are dysregulated at a protein level in Fmr1 KO astrocytes, and that secretory machinery including clathrins and Golgi adaptor proteins are downregulated in Fmr1 KO. Suppressing BMP signaling with Smad4 cKO improves this Fmr1 KO deficit in secretory machinery at the level of the entire pathway. BMP signaling thus could control bulk level of astrocyte-secreted proteins in Fmr1 KO.^12^ BMP signaling’s effect on vesicular machinery could also affect secretion of other compounds not detectable by ER-TurboID, such as peptides, lipids, and proteins without external lysine residues to enable proximity labeling.^58,90^ Applying this ER-passaged astrocyte proteomic approach to other genetic neurodevelopmental disorder models will allow investigation of whether there are convergent mechanisms of *in vivo* astrocyte secretome dysregulation across disorders, as was observed *in vitro*^12^ and would be predicted by a convergent neuroscience approach to autism spectrum disorder.^91^

Previous studies have shown mice globally heterozygous for the BMP receptor Bmpr2 have rescued Fmr1 KO spine deficits,^14^ cultured Fmr1 KO astrocytes have an elevated BMP signaling transcriptional signature,^12^ and P7 cortical Fmr1 KO astrocytes have elevated canonical BMP signaling.^12^ Cell type-specific bulk RNAseq and scRNAseq in P7 to P14 cortex show ongoing astrocyte BMP signaling capability (*Bmpr1b* enriched in astrocytes) and effect (*Id3* enriched in astrocytes, vascular/leptomeningeal cell, and endothelial cells).^53,92^ Surprisingly, in our astrocyte nuclear RNAseq at P26-28, we did not detect elevated expression of *Id* genes (BMP-induced) or *Smad6* (expected BMP pathway negative feedback inhibition)^15^ in Fmr1 KO (Comparison 1). However, further detailed examination of BMP-pathway related genes shows in Fmr1 KO (Comparison 1) a trending increase in the ligand *Bmp7* (L2FC = 0.60, p-adj = 0.11) which is restored to WT levels in FXS;Smad4 cKO, as well as a significant increase in expression of Chrdl1 (WT to FXS L2FC = 0.75, p-adj = 0.027), a secreted protein that binds to and inhibits BMPs (**Supplementary Data Table 5**).^93^ One hypothesis is that by P28, earlier elevations in astrocyte BMP signaling led to compensatory negative feedback such as upregulation of the inhibitor Chrdl1, resulting in normalized signaling. Within this model, earlier upregulation of astrocyte BMP signaling may still contribute to dysregulated formation of neuronal synapses and circuits. Moreover, the Smad4 cKO intervention, driven by a Cre driver line characterized to be active by P0-P7,^29^ could still counteract a temporally restricted BMP signaling increase.

We found that suppressing canonical astrocyte BMP signaling with Smad4 cKO lessens Fmr1 KO audiogenic seizure severity and rescues Fmr1 KO inhibitory synapse deficits in auditory cortex. The decrease in Fmr1 KO auditory cortex inhibitory synapses we see is similar in size to that observed by others,^76^ and in line with cortical hyperexcitability observed across multiple Fmr1 KO cortical regions.^78^ The restoration of inhibitory synapses in auditory cortex by Smad4 cKO is one mechanism that may reduce auditory hypersensitivity. Importantly, the Smad4 cKO reduction in Fmr1 KO seizure severity was not a complete rescue to the WT level. This moderating effect may reflect the partial KO (∼36±3% overall, 61±4% superficially) of astrocyte SMAD4 in inferior colliculus (**Supplementary Fig. 1E,F**), a region linked to audiogenic seizure etiology.^94^ Importantly, Smad4 cKO did not improve either of the two visual phenotypes tested, suggesting mechanisms independent of astrocyte BMP signaling may underlie Fmr1 KO visual acuity and ocular dominance plasticity phenotypes.

What pathways specifically mediate Smad4 cKO improvements to Fmr1 KO phenotypes? Our transcriptomic and proteomic analyses find that astrocyte metabolic and secretory pathways are dysregulated in Fmr1 KO and improved when BMP signaling is suppressed; these are promising candidate pathways for mediating the Smad4 cKO effect. Additionally while hepaCAM, a regulator of inhibitory synaptogenesis, is not changed in Fmr1 KO, Smad4 cKO leads to an increase in hepaCAM that could mitigate the Fmr1 KO baseline deficit in inhibitory synapses (**Supplementary Fig. 11D**). Alternatively, Smad4 cKO may act through the aggregated and/or interacting effects of multiple downstream actors, as could be expected given its role as a transcriptional regulator known to target hundreds of genes.^26^ As secretory machinery as a whole is downregulated in Fmr1 KO, by improving the Fmr1 KO deficit in secretory machinery, Smad4 cKO may help restore the levels of many secreted products. This may be analogous to what was observed *in vitro*, where astrocyte BMP pathway suppression but not individual downstream protein targeting rescued Fmr1 KO neurite outgrowth.^12^ Thus, whether through specific proteins, improvements to astrocyte metabolism or secretion, or a combination of the above mechanisms, suppressing the BMP signaling pathway in astrocytes with Smad4 cKO can improve Fmr1 KO phenotypes.

Taken together, this study sheds light on the role of astrocyte BMP signaling in FXS phenotypes and provides leads on several astrocyte pathways that mechanistically connect BMP signaling to Fmr1 KO phenotype correction and thus represent candidates for targeted interventions.

## Methods

### Mice

All animal work was approved by the Institutional Animal Care and Use Committee (IACUC) of the Salk Institute for Biological Studies. Mice were housed in the Salk Institute Animal Resources Department on a light cycle of 12 hours light:12 hours dark with access to food and water *ad libitum*. For experiments using developmental timepoints, littermates were used.

### Fmr1 KO mice

B6.129P2-Fmr1tm1Cgr/J (Jax: 003025) line Fmr1 WT and KO male mice were used for experiments. We used male mice because female fragile X syndrome (FXS) patients (and mice) are heterozygous and mosaic for the *FMR1* gene due to random X inactivation, leading to less severe and more variable clinical manifestations.^36^ Hemizygous male to heterozygous female crosses were used. The line was maintained on a C57Bl6/J background.

### Fmr1 KO x Smad4 fl/fl x Gfap-Cre mice

The Fmr1 KO line was crossed to Smad4tm2.1Cxd/J (Smad4 fl/fl, Jax: 017462) and B6.Cg-Tg(Gfap-Cre)73.12Mvs/J (Jax: 012886) lines to remove Smad4 from astrocytes (Smad4 cKO). Experimental mice were obtained from three types of breedings: (1) Fmr1 WT;Smad4 fl/fl male x Fmr1 WT;Smad4 fl/fl;Gfap-Cre+ females to generate Fmr1 WT;Smad4 WT (referred to as WT) and Fmr1 WT;Smad4 cKO (referred to as WT;Smad4 cKO) male littermate pairs (2) Fmr1 WT;Smad4 fl/fl male x Fmr1 KO;Smad4 fl/fl;Gfap-Cre+ females to generate Fmr1 KO;Smad4 WT (referred to as FXS) and Fmr1 KO;Smad4 cKO (referred to as FXS;Smad4 cKO) male littermate pairs and (3) Fmr1 KO;Smad4 fl/fl male x Fmr1 het;Smad4 fl/fl;Gfap-Cre+ female to generate all four of the experimental genotypes as well as Fmr1 KO females for line maintenance. The line was maintained on a C57Bl6/J background.

### Cell cultures

#### Primary astrocyte cultures

Astrocytes were obtained from the cerebral cortex of P0-2 C57Bl6/J background mice following the protocol described in McCarthy and de Vellis, 1980^95^ and Blanco-Suarez et al., 2018.^39^ Cortices of 4 mice were dissociated using papain (Worthington PAP2 3176) and plated on 75cm2 tissue culture flasks coated with poly-D-lysine (Sigma P6407), then maintained in culture for 4 days in growth medium (DMEM [Thermo Fisher Scientific 11960044] supplemented with 10% heat inactivated FBS [Thermo Fisher Scientific 10437028], Penicillin-Streptomycin [Thermo Fisher Scientific 15140-122], glutamax [Thermo Fisher Scientific 35050-061], sodium pyruvate [Thermo Fisher Scientific 11360-070], hydrocortisone [Sigma H0888], and N-acetyl-L-cysteine [Sigma A8199]). Astrocytes form a monolayer adherent to the flask bottom, and after 3 days, flasks were shaken vigorously to remove non-adherent cells on overgrown layers. Two days later AraC (working concentration 10 mM; Sigma C1768) was added for 48 hr in order to inhibit proliferation and kill rapidly proliferating cells such as oligodendrocyte precursor cells and microglia. After two more days, astrocyte-enriched cells were dissociated from the flask with trypsin, then plated on 10 or 15 cm tissue culture dishes and maintained in the growth media described above. Cells were passaged once a week, and cultures were maintained for no longer than 4 weeks. Astrocyte cultures were maintained in a humidified incubator at 37C and 10% CO2.

### Method Details

#### Tissue collection

Mice were deeply anesthetized with intraperitoneal injection of a mixture of 100mg/kg ketamine (Victor Medical Company 1699053) and 20mg/kg xylazine (Victor Medical Company 1264078). For acute astrocyte isolation and for fresh tissue collection for transcriptomics, proteomics, and western blotting experiments, cortices were dissected immediately following deep anesthesia and cervical dislocation. For fixed tissue collection for histological experiments, anesthetized mice were transcardially perfused first with room temperature PBS followed by 4% PFA. The brains were rapidly dissected and post-fixed overnight in 4% PFA (Electron Microscopy Sciences 19210) at 4C. After 16-24 hours the brains were moved to 30% sucrose for 48-72 hours. The brains were then embedded in TFM (Electron Microscopy Sciences 72593) and frozen in a dry ice/ethanol slurry and stored at -80C until use.

#### Acute astrocyte isolation

Astrocytes were acutely isolated from P14 mouse cortex with a protocol modified from that described in Brandebura et al., 2023.^45^ Dissected cortices were cut into ∼1mm^3^ pieces using a razor blade to allow for optimal enzymatic digestion. Papain digestion was performed according to kit instructions (Miltenyi MACS Neural Tissue Dissociation Kit 130-092-628). Digestion was performed using the gentleMACS Octo Dissociator with Heaters (Miltenyi 130-096-427) using the 37C_NTDK program. Cells were centrifuged at 300 g 4C for 10 minutes and then resuspended in AstroMACS buffer and passed through a 70 um basket filter to remove clumps. Astrocytes were subsequently isolated using the Miltenyi MACS Anti-ACSA-2 Microbead kit (Miltenyi 130-097-678) according to the manufacturer’s instructions, with modifications to deplete debris and microglial cells prior to positive astrocyte selection. Debris was removed with the Debris Removal Solution (Miltenyi 130-109-398), following manufacturer instructions, then the depleted solution was diluted to 15mL in cold DPBS and centrifuged at 1000 g 4C for 10 minutes. Microglia depletion was performed using anti-CD11B beads (Miltenyi 130-093-634) for an incubation period of 10 minutes at 4C. The flow-through from the MS magnetic column was collected and centrifuged again at 300 g 4C for 10 min to remove excess beads. The pellet was collected and subsequent positive selection for astrocytes was performed. First a 10-minute incubation at 4C with FcR blocking beads was used to minimize any nonspecific binding. ACSA2-conjugated beads were then used to positively select for astrocytes during a 15-minute incubation period at 4C (FcR and ACSA2 beads in kit, Miltenyi 130-097-678). Bound astrocytes were plunged into a new tube with AstroMACS buffer and centrifuged again at 300 g 4C for 10 min to remove excess beads. The astrocytes were divided and frozen at -80C either in lysis solution (for subsequent Western blotting; Pierce RIPA [Thermo 89901] with 1:100 phosphatase [Thermo 1862495] and protease [Thermo 78429] inhibitors) or as a pellet (for subsequent DNA extraction).

#### Western blotting

Western blotting was performed on acutely isolated astrocytes to validate Smad4 cKO, on proteomics samples to validate biotinylation, and on cortical lysates to validate brevican proteomics data. Acute astrocytes frozen in lysis buffer were thawed, homogenized with pellet mixer (VWR 47747-370), and spun down (16800 rpm, 20 min, 4C). For cortical lysates, one cortex was homogenized with the pellet mixer in lysis buffer (Pierce RIPA with 1:100 phosphatase and protease inhibitors), allowed to rotate for 1hr at 4C, then spun down (20000 g, 15 min, 4C). The concentration of the supernatant was quantified with BCA (Thermo 23227), and 10ug acute astrocyte protein or 20ug cortical lysate was used per sample.

Samples were prepared with reducing sample buffer (Thermo Fisher Scientific 39000) and boiled for 10 min at 95C (acute astrocytes and lysates for ER-TurboID validation) or 45 min at 55C (cortical lysates). Proteins were separated on Bolt 12% Bis-Tris Plus gels (NW00120BOX) for 1 hr at constant voltage of 130V in Bolt MOPS SDS running buffer (Thermo B0001_02). Proteins were transferred onto Immobilion-FL PVDF membranes (Millipore IPFL00010) at 100V for 60 min for acute astrocytes, or onto Amersham Hybond 0.2 um PVDF membranes (Cytiva 10-6000-22) at 80V for 120 min in a solution of Tris-Glycine Buffer (Thermo Fisher Scientific 28363). Membranes were blocked for 1 hr at room temperature in gentle shaking with blocking buffer (0.1% Casein [BioRad 1610782] in Tris Buffered Saline [TBS]). Membranes were incubated overnight at 4C with gentle shaking with the primary antibody in blocking buffer, rabbit anti-SMAD4 1:1000 (Millipore ABE21), mouse anti-BCAN 1:4000 (BD Biosciences 610894), mouse anti-beta-Actin 1:5000 (Millipore Sigma A1978), mouse anti-TUBB3 1:5000 (BioLegend 801201), rabbit anti-Actin 1:5000 (Cell Signaling Technology 4968), or rabbit anti-SOX9 1:2000 (Abcam AB185966). Membranes were washed with TBS+0.1% Tween and incubated for 1 hr at room temperature with secondary antibody goat anti-mouse Alexa 680 (1:10000, Thermo Fisher Scientific A21058), goat anti-rabbit Alexa 680 (1:10000, Thermo Fisher Scientific A21109), and goat anti-mouse DyLight 800 (1:10000, Invitrogen SA535521). For ER-TurboID validation westerns, no primary step was performed, and membrane blocking was followed immediately by washes and secondary (1:3000, IRDye 680RD Streptavidin, Li-Cor 926-68079). Membranes were washed with TBS+0.1% Tween, and immediately imaged on Odyssey Infrared Imager CLx (Li-Cor) using Image Studio v5.2 software (Li-Cor). Molecular weights were visualized using Precision Plus All Blue Protein standard loading dye (BioRad 1610373) or PageRuler (Thermo 26616). Analysis of band signal intensity was done in Image Studio v5.2 software (Li-Cor). For SMAD4, signal was normalized to SOX9 intensity in different lanes the same blot (cut membrane); for BCAN, signal was normalized to actin intensity in the same lane of the same blot (cut membrane).

#### DNA extraction and PCR

For validation of Smad4 cKO recombination at the Smad4 exon 9 locus, DNA was isolated from acutely isolated astrocytes and PCR for the recombined product was performed. DNA was isolated with the DNeasy blood and tissue kit (Qiagen 69504), following manufacturer instructions. PCR was performed using SYBR green PCR mastermix (Life Supply 4309155), and the below primers from those used by the creators of the Smad4 fl/fl line^27^ with modifications to align with the mm10 genome:

Forward (5’ to 3’): TAA GAG CCA CAG GGT CAA GC

Reverse (5’ to 3’): GAC CCA AAC GTC ACC TTC AC

PCR products were run on a 1.5% agarose gel with expected band size of 500bp in the recombined sequence and no band expected in WT.

#### Immunohistochemistry and analysis: Smad4 cKO validation

The cell type specificity of Smad4 cKO was evaluated with immunohistochemistry (IHC) for SMAD4, SOX9, and NEUN. Validation of suppression of BMP signaling in astrocytes was performed by IHC for ID3 and SOX9. Sagittal brain sections of 16 um were obtained from littermate WT and WT;Smad4 cKO mice (1.8mm lateral from midline) using a cryostat (Hacker Industries OTF5000). Sections were on Superfrost Plus micro slides (VWR 48311-703) for this as well as all other experiments involving cryosectioning. Slices were blocked and permeabilized in blocking buffer (10% donkey serum [Jackson ImmunoResearch 017-000-121] in PBS with 0.3% Triton [Sigma T9284] and 100mM L-lysine [Sigma L5501]) for one hour at room temperature. Primary antibodies were diluted in antibody dilution solution (5% donkey serum in PBS with 0.3% Triton and 100mM L-lysine) and stained overnight in a humidified chamber at 4C. Antibodies for the astrocyte marker SOX9 (1:100, goat, R&D AF3075), neuronal marker NEUN (1:200, mouse, EMD Millipore MAB377), SMAD4 (1:250, rabbit, Millipore ABE21), and ID3 (1:200, rabbit, CST9837S) were used. The next day sections were washed three times in PBS with 0.2% Triton at 5, 10, and 15 minutes. Secondary antibody labeling was performed in antibody solution (5% donkey serum in PBS with 0.3% Triton and 100mM L-lysine) and for two hours at room temperature. Appropriate secondary antibodies were used (1:500, donkey anti-goat Alexa 488, Jackson ImmunoResearch 705-545-147; 1:500, donkey anti-rabbit Alexa 568, Invitrogen A10042; 1:500, donkey anti-mouse Alexa 647, Jackson ImmunoResearch 715-605-151). Washes in PBS with 0.2% Triton were performed three times at 5, 10, and 15 minutes. DAPI was applied (1:1000, Thermo 62247) for 5 minutes followed by a PBS wash. Coverslips (Fisher 12541037) were mounted with SlowFADE Gold Antifade Mountant with DAPI (Invitrogen S36939).

SMAD4 validation IHC was imaged on an LSM700 confocal with 40x oil-immersion objective (numerical aperture [NA] = 1.3) as 16-bit images, as a z-stack of 8 steps of 0.4 um each spanning 2.8 um. Images were collected as tiles (12 tiles for cortex, consisting of 6143 x 2047 pixels spanning L1 to L5-L6, 4 tiles for colliculi and medial geniculate nucleus [MGN] of the thalamus, consisting of 2047 x 2047 pixels, all with 0.16 um x 0.16 um pixel size, pixel dwell time 1.58 us). Analysis was performed using Imaris software (Bitplane) and the ImageJ LabKit plug in to automate detection of SOX9+ and NEUN+ nuclei, within which SMAD4 fluorescence intensity was quantified. Within each experiment (littermate pair), the combined distribution of nuclei SMAD4 intensities was used to assign a single threshold for SMAD4 positivity at a local minimum in the SMAD4 intensity distribution. Cells below the threshold were visually inspected and compared to surrounding parenchymal intensity to confirm lack of SMAD4 positivity.

ID3 IHC was imaged on an AxioImager Z.2 with Apotome module with 20x objective (NA = 0.8) as 14-bit images as 6 tiles spanning all layers of cortex and white matter (3887 x 1975 pixels, 0.323 um x 0.323 um per pixel), as a z-stack of 4 steps of 2 um each with total thickness 6 um. Analysis was performed using Imaris software (Bitplane) with the surfaces tool to automate detection of SOX9+ nuclei with parameters: smoothing 0.300 um, diameter 10 um, threshold intensity 20, volume filtered above 150 um^3^, sphericity filtered above 0.5. Within SOX9+ nuclei, ID3 intensity was quantified, with layer stratification performed using DAPI staining. Images shown in **Fig. 1C** and **Supplementary Fig. 1E,G** are maximum intensity projections.

#### Immunohistochemistry and analysis: Smad4 cKO model characterization

To quantify cortical density of S100B+ cells, immunohistochemistry was performed as above on sagittal slices from Smad4 WT and Smad4 cKO littermate pairs, with primary antibody astrocyte marker S100B (1:500, rabbit, Abcam 52642). Following imaging under the same parameters as for SMAD4 IHC above, S100B density was quantified with ImageJ using the cell counter plug-in and counting only S100B cell bodies colocalized with DAPI. Manual separation of layers was performed with visualization by DAPI. DAPI density was quantified with Imaris software (Bitplane), with automated surface creation with parameters: smoothing 0.313 um, diameter 5 um, threshold intensity 30, volume filtered 50 to 750 um^3^, automatic seeding with predicted distance 5 um. Separation by layer was performed using DAPI staining. Images shown in **Supplementary Fig. 3A,B** are maximum intensity projections.

#### Behavioral analysis

Behavioral testing was performed in the facilities of Salk Institute’s In Vivo Scientific Services (Animal Resources Department). All behavioral tests were performed blinded to genotype.

#### Audiogenic seizure assay

The audiogenic seizure assay was performed at a timepoint within P20-P23, when Fmr1 KO mice are reported to have high susceptibility.^96^ Mice were transported to the IVSS behavior core and allowed 30 min to adapt to the low level of ambient noise (∼40dB). Mice were then individually taken to a separate soundproofed room and placed in a sound-proofed chamber equipped with light, video camera, plexiglass box (7.5 x 11.5 x 5 inches), and alarm (115-125dB, Kosin Safe Sound Personal Alarm ASIN#B07NLFQS7T). After a 5-minute habituation period, the alarm was played for 2 minutes, with video recording throughout. Each mouse would then be scored numerically as 0 (no seizure activity), 1 (convulsive activity), and 2 (respiratory failure) for the maximum degree of severity reached during the 2-minute period. Others have reported a scale of 0-3 including a stage for wild running that precedes convulsive activity;^33^ in our data, we found every mouse that had wild running progressed further to convulsive activity. Testing was always performed on all littermates in one session, blind to genotype.

#### Optomotor assay

The optomotor assay was performed at P26 with a setup from CerebralMechanics (2004) (elevated platform 5 cm above top and bottom mirrors with drifting gratings at 12 deg/sec created through four computer monitors encircling the platform) and software from Optomotry VR, with settings as staircase method from 0.042 cycles/deg to 0.5 cycles/deg, scoring both clockwise and counter-clockwise visual acuity. The average of these two readings was the reported visual acuity value for all data and figures in this manuscript. This setup and procedure is described in further detail by Prusky et al., 2004.^37^ Mice were always tested in Fmr1 WT/KO or Cre +/-pairs, blind to genotype.

#### Monocular visual enucleation and *Arc* analysis

Littermate WT and Fmr1 KO mice as well as littermate FXS and FXS;Smad4 cKO mice at P28 (within visual critical period) were used for these experiments, which were performed as described previously.^39^ Mice were anesthetized with 2% isofluorane in oxygen by constant flow via nose cone. The right eye was removed by transecting the optic nerve. Gelfoam (Pfizer 031508) was placed in the empty ocular cavity. Eyelids were sutured using 6-0 silk sutures (Henry Schein 101-2636). Lidocaine 2% and erythromycin 0.5% were applied topically on the sutured eyelids. Normal rearing (NR) mice were collected 12 hr later. Mice for monocular enucleation (ME) analysis were subjected to a 12 hr light/12 hr dark cycle every day until collection 4 days later, at P32. Prior to collection, mice were exposed to light for 30 min in alert condition to activate neurons in the visual cortex and induce expression of *Arc*. Mice were anesthetized with a mixture of 100mg/kg ketamine (Victor Medical Company 1699053) and 20mg/kg xylazine (Victor Medical Company 1264078) and brains removed. Brains were coronally sectioned in half, and the posterior brain collected and embedded in OCT (Scigen 4583), frozen in dry ice/ethanol, and stored at -80C.

#### Single molecule fluorescent *in situ* hybridization (smFISH)

Littermate WT and Fmr1 KO male mice as well as littermate FXS and FXS;Smad4 cKO male mice that underwent monocular enucleation were used to analyze *Arc* expression to characterize ocular dominance plasticity, with modifications from protocols described previously.^39^ All sections were made using a cryostat (Hacker Industries OTF5000) at a slice thickness of 16 um, 3.40 mm posterior to bregma for coronal sectioning. Fluorescent *in situ* hybridizations (FISH) were performed with RNAscope kits (ACDBio 320851 for V1, used for Fmr1 KO vs. WT; ACDBio 323100 for V2 used for FXS vs. FXS;Smad4 cKO) following manufacturer’s instructions for fresh frozen tissue, with the following modifications. For V1, tissue was treated with Protease 4 for 12 min at room temperature, and *Arc* probe (ACDBio 316911) was conjugated to the 550 channel with AltB, and SlowFade gold antifade mountant with DAPI was applied to every section. For V2, tissue was treated according to manufacturer instructions with the following modifications. Slides were fixed by immersing in prechilled 4% PFA in PBS for 60 min at 4C, after dehydration slides were baked at 37C for 30 min, Protease III was applied for 15 min at room temperature, *Arc* probe was diluted by 50% with ACDBio probe diluent (ACDBio 300041), and Opal fluorophore 570 (Akoya FP1488001KT) was used at 1:5000 dilution. Coverslips 22 mm X 50 mm 1.5 thickness were placed on top of the sections and sealed with nail polish (Sally Hansen 45111). In control experiments a negative control probe was used (ACDBio 310043) to determine the level of background signal. For *Arc* visualization, coronal sections were imaged using fluorescence microscope Zeiss AxioImager.Z2 at 10X magnification as 14-bit images of 100-200 tiles. *Arc* was imaged in the 550 channel. Zen Blue software distance tool was used to measure the width of the *Arc* activated brain region along layer 4 of the visual cortex, contralateral to the removed eye. 2-8 brain sections were measured per mouse, and a minimum of 4 animals were used per condition. Analysis was performed blind to genotype.

#### Glyoxal-fixed astrocyte nuclei transcriptomics

##### Sample preparation

Samples for transcriptomics were prepared as in Labarta-Bajo et al., 2023.^41^ Each sample consisted of two cortices from one mouse; mice were processed from tissue collection to fluorescence-activated cell sorting (FACS) in littermate Cre+/-pairs, then nuclear lysates were stored in Buffer RLT (Qiagen 79216) at -80C. All samples were then thawed and processed in parallel from RNA extraction to sequencing. Samples were from the four genotypes WT, WT;Smad4 cKO, FXS, FXS;Smad4 cKO, with 4 to 6 mice per genotype as biological replicates.

##### Nuclei preparation

Nuclei were isolated from freshly collected cortex of P26-28 mice. Cortices were dissected into fragments approximately 9 mm^3^ in size and placed in glyoxal acidic solution (3% w/v glyoxal [Millipore Sigma 128465], 20% ethanol [Fisher BP2818100], 0.75% acetic acid [Millipore Sigma A6283], 5mM NaOH [Millipore Sigma 72068], in ddH2O) for 15 min at 4C. Tissue was manually homogenized using a two-step Dounce homogenizer (A and B) (Sigma #D9063) in NIMT buffer (in mM: 250 sucrose, 25 KCl, 5 MgCl2, 10 Tris-Cl pH 8, 1 DTT; 1:100 dilution of 10% Triton and protease inhibitor [Halt 78429]) on ice. Homogenized samples were mixed with 50% iodixanol (OptiPrep Density Gradient Medium; Sigma #D1556) and loaded onto 25% iodixanol cushion, and centrifuged at 10000 g for 20 min at 4C in a swinging bucket rotor (Sorval HS-4). Pellets resuspended in ice-cold DPBS with 1% w/v BSA (Millipore A7030) (referred to as DPBS+BSA). Nuclei were then incubated for 7 min on ice with Hoechst 33342 solution (2 mM) (Thermo #62249) (final concentration 5 μM), followed by centrifugation at 1000 g for 5 min at 4C to pellet nuclei. Pellets were resuspended in blocking buffer containing DPBS+BSA with NEUN-Alexa 488 pre-conjugated antibody (Millipore #MAB377X) at 1:1000 dilution along with SOX9 rabbit antibody (Abcam AB185966) at 1:200, and incubated for one hour at 4C. Following centrifugation at 1000 g for 5 min at 4C to pellet nuclei, pellets were resuspended in DPBS+BSA with goat anti-rabbit-Alexa 647 (Invitrogen A21245) at 1:200 dilution for one hour at 4C. Tubes were centrifuged at 1000 g for 5 min at 4C then resuspended in 500ul DPBS+BSA and filtered with a 100 um cell strainer (Corning 431752).

##### Flow cytometry and RNA extraction

FACS was performed in the Salk Institute Flow Cytometry core using a BD FACS Aria Fusion sorter with PBS for sheath fluid (a 70 um nozzle was used for these experiments). Hoechst-positive nuclei were gated first (fluorescence measured in the BV421 channel), followed by exclusion of debris using forward and side scatter pulse area parameters (FSC-A and SSC-A), exclusion of aggregates using pulse width (FSC-W and SSC-W), before gating populations based on SOX9 and NEUN fluorescence (using the APC and FITC channels). To isolate the astrocyte population, nuclei devoid of FITC signal (NEUN-) but with APC signal (SOX9+) were collected (**Supplementary Fig. 5A**). Nuclei were purified using a one-drop single-cell sort mode (for counting accuracy); these were directly deposited into a 1.5ml Eppendorf with 300ul DPBS+BSA preloaded. Approximately 200,000 nuclei were collected per mouse (range: 115k to 276k). After sorting each mouse, the four collected samples (SOX9+NEUN-, SOX9-NEUN-, SOX9+NEUN+, SOX9-NEUN+) were each checked for purity using the same gates. Illustrations of gating strategy and post-sort purity were created with FlowJo v10.10.0. Immediately after sorting, nuclei were centrifuged at 1000 g for 5 min at 4C, then resuspended in RLT buffer with added β-mercaptoethanol (Sigma M3148), then frozen at -80C.

RNA was extracted from FACS-sorted nuclei using the RNeasy micro kit (QIAgen 74104). The homogenate was thawed at room temperature then RNA was extracted according to manufacturer instructions, with on-column DNase digestion (Qiagen 74004) and elution in 14ul water at 65C.

##### RNAseq

RNA sequencing libraries were generated using the Illumina Ribo-Zero Plus rRNA Depletion Kit with IDT for Illumina RNA UD Indexes (Illumina, San Diego, CA, #20040525). Samples were processed following manufacturer’s instructions. Average input RNA per sample as measured by TapeStation was 79ng (range: 26-203ng). Samples with TapeStation traces indicating degraded RNA were not sequenced. Resulting libraries were multiplexed and sequenced with 100 basepair (bp) Paired End (PE100) to an average depth of 25 million reads per sample (range of 19 million to 39 million reads) on an Illumina NovaSeq 6000. Samples were demuxltiplexed using bcl2fastq v2.20 Conversion Software (Illumina, San Diego, CA).

##### RNAseq analysis

RNAseq sequencing data was mapped with STAR v2.5.3a^97^ to the mm10 genome, and FASTQ and STAR alignment files were quality tested with MultiQC v1.0.dev0^98^ (average uniquely mapped reads 87.7%, range 85.1% to 91.2%). Genes were called with Homer v4.10.4^99^ (range 17 to 35 million mapped tags per sample). Differential analysis was conducted with DESeq2 (v1.38.3),^100^ with differentially expressed gene criteria as follows: average TPM > 1 in WT samples, Benjamini-Hochberg (BH) adjusted p-value < 0.05, |fold change (FC)| > 1.5. Heatmaps were created using R (v4.2.2) via RStudio (2022.12.0) using the Pheatmap package and Graphpad Prism (v8.0). Volcano plots were created with package ggplot2 (v3.4.0). Gene set enrichment analysis (GSEA) was performed to detect which pathways and gene sets were enriched in Fmr1 KO and Smad4 cKO using the package ClusterProfiler (v4.6.0).^101^ The following gene sets from mouse MSigDB^102^ were used: hallmark pathways, canonical pathways (including Reactome,^103^ KEGG,^104^ Wikipathways,^105^ and BioCarta^106^), and gene ontology.^107,108^ Genes with TPM > 1 in every sample were used for GSEA, with ranking based on sign(L2FC)*-log10(p-value), for each comparison. Highlighted pathways in visualization are top three down- and upregulated (lowest and highest normalized enrichment score [NES]) hallmark pathways in each comparison (**Fig. 3E**). For testing for overlap of known FMRP targets with up- and down-regulated genes in Fmr1 KO (Comparison 1), the list of 842 genes from Darnell et al. 2011^52^ was used, and statistics were done with Fisher’s exact test with genome size 24,252 protein-coding *M. Musculus* genes. For testing of the distribution of L2FC values for all genes in Fmr1 KO nRNAseq (20,482 detected) compared to L2FC values for genes known to be FMRP targets^52^ and enriched in astrocytes^53^ (141 genes), the z-test was used (**Fig. 2E**).

#### ER-TurboID proteomics

##### Cloning of plasmids for viral vectors

The pAAV-TBG-Cyto-TurboID, pAAV-TBG-ER-TurboID, Sec61b-V5-TurboID, and C1(1-29)-TurboID-V5_pCDNA3 vectors were obtained from Addgene (149414, 149415, 166971, 107173).^58–60^ Each was separately inserted into the pZac2.1 GfaABC1D-tdTomato vector^109^ obtained from Addgene (44332) after removing tdTomato sequence. The In-Fusion HD cloning kit (Takara 638909) was used according to the manufacturer’s instructions, with analysis in SnapGene v6.1.2. The GfaABC1D-tdTom vector was linearized at NheI and NotI (New England Biolabs R3131S, R1089S) restriction sites for three hours at 37C and the linearized vector was purified from an agarose gel and cleaned using Nucleospin columns (Macherey-Nagel 740609.50). The C1(1-29)-TurboID, Cyto-TurboID, ER-TurboID, and Sec61b-V5-TurboID cDNA were PCR amplified.

For the Cyto-TurboID, a cytosine was added before the start codon, and an HA tag and Ala Ser linker were added before the TurboID sequence, using the below primer sequences with 15-18bp overlapping sequences to the linearized vector:

Forward (5’ to 3’): CTC ACT ATA GGC TAG CGC CGC CAT GTA TCC GTA TGA TGT TCC GGA TTA TGC AGC TAG CAA AGA CAA TAC TGT GCC T

Reverse (5’ to 3’): GCT CGA AGC GGC CGC TTA GTC CAG GGT CAG GCG CTC CAG GG

For the ER-TurboID, the below primer sequences with 17bp overlapping sequences to the linearized vector were used:

Forward (5’ to 3’): ACT CAC TAT AGG CTA GCG CCA CCA TGG AGA CAG AC

Reverse (5’ to 3’): CTG CTC GAA GCG GCC GCT TAG AGT TCA TCT TTG GTG CTG TCC AG

For Sec61-V5-TurboID, GCCACC was added before the Sec61b sequence and Ala Ser and an HA tag were added before the stop codon, using the below primer sequences with 15-18bp overlapping sequences to the linearized vector:

Forward (5’ to 3’): TCA CTA TAG GCT AGC GCC ACC ATG CCT GGT CCG ACC CC Reverse (5’ to 3’): CTC GAA GCG GCC GCT TAT GCA TAA TCC GGA ACA TCA TAC GGA TAG CTA GCC TTT TCG GCA GAC CGC A

The PCR product was subsequently gel purified and cleaned with the Nucleospin columns. The In-Fusion HD cloning enzyme premix was used to insert the above four amplified sequences into the vector.

<u>The C1(1-29)-TurboID</u> was cloned into the GfaABC1D-tdTom vector using the same procedure, but with KpnI HF and XhoI restriction enzymes (R3142S, R0146S). The primer sequences used are as follows:

Forward (5’ to 3’): CTC ACT ATA GGC TAG CGC CAC CAT GGA Reverse (5’ to 3’): CTC GAA GCG GCC GCC TAT GCA TAA TC

Stellar competent *E. Coli* cells (Takara 636763) were then transformed with the plasmids via heat-shock according to the manufacturer’s instructions. Bacteria were screened using carbenicillin resistance (100 ug/mL; Teknova C2130). The colonies were expanded in an overnight LB culture. The DNA was purified using an endotoxin-free plasmid MaxiPrep kit according to the manufacturer’s instructions (Qiagen 12362). Newly created constructs were sequenced with Eton sequencing (all constructs) as well as Primordium labs (ER-TurboID) to confirm desired product prior to following experiments.

##### Testing ER-TurboID constructs in astrocyte cultures

Five days after passaging into a 10cm plate, the mouse astrocytes were transfected with OptiMEM (Thermo 31985062, 1800ul), lipofectamine 2000 (Thermo 11668-019, 45ul), and 15ug plasmid: either control CAG-GFP, GfaABC1D-C1(1-29)-TurboID, GfaABC1D-cyto-TurboID, GfaABC1D-ER-TurboID, or GfaABC1D-sec61b-TurboID. Three hours after the start of transfection, astrocytes were switched to high protein conditioning media without biotin (components 50% Neurobasal Thermo 12348017, 50% DMEM LifeTech 11960044, supplemented with Penicillin-Streptomycin [Thermo Fisher Scientific 15140-122], glutamax [Thermo Fisher Scientific 35050-061], sodium pyruvate [Thermo Fisher Scientific 11360-070], insulin [Sigma I6634], N-acetyl cysteine [Sigma A8199], transferrin [Sigma T-1147], BSA [sigma A4161], progesterone [Sigma P8783], putrescine [Sigma P5780], sodium selenite [Sigma S9133], and T3 [Sigma T6397]). 24 hours after transfection, biotin (Sigma B4639) was added to 50uM in the media. After 17 hours, the media was changed to low protein conditioning media without biotin for 21 hours, after which lysates were collected by adding 200ul RIPA lysis buffer (Thermo 89901) with 1:100 protease (Thermo 78429) and phosphatase (Thermo 1862495) inhibitors directly to the cells and applying a cell scraper (Thermo 179693) to ensure lysis. Protein concentration was quantified with BCA kit (Thermo 23227), then 75ug protein from each lysate sample was added to 15ul streptavidin beads (Thermo 88817), incubated for 1hr at room temperature and 16hr 4C rotating, washed at room temperature with 250ul each of RIPA lysis buffer (2x2 min), 1M KCl (Sigma P9591) (1x2 min), 0.1M sodium carbonate (Sigma 222321) (1x10 s), 2M urea (Invitrogen 15505-035) in 10mM Tris-HCl pH 8 (1x10 s), RIPA lysis buffer (2x2 min, moving to a new tube), and eluted in 30ul elution buffer (protein loading buffer [Thermo 39001] diluted to 1x, 20mM DTT [MilliporeSigma D9779], and 2mM biotin [Sigma B4639]) at 95C for 10 min. Washes were done and eluates were collected with magnetization of beads with a magnetic stand (DynaMag-2 Magnet, Thermo 12321D). Streptavidin western blots were performed as described in the Western blotting section above.

##### Viral vector generation

Plasmids were packaged into AAV at the Salk Institute Viral Vector (GT3) core facility. Viruses were packaged as AAV PHP.eB serotype with titer of approximately 5.6E14 genome copies (gc)/mL (GfaABC1D-ER-TurboID, plasmid for biotinylating enzyme) and 2.38E14 gc/mL (GfaABC1D-smFP, plasmid for control protein, subcloned as described in Brandebura et al. 2023,^45^ based on construct from Viswanathan et al., 2015^110^).

##### Retroorbital virus injections and biotin supplementation

Mice at P12 were anesthetized with 2-3% isofluorane in oxygen by constant flow through a nose cone. Virus was injected via retroorbital administration at a dose of 1.0E12 viral genome copies in a total injection volume of 90ul with either AAV-PHP.eB-GfaABC1D-ER-TurboID or AAV-PHP.eB-GfaABC1D-smFP. The virus was diluted in DPBS just prior to injection. Mice were injected with subcutaneous biotin (24mg/kg, 5mM biotin in PBS) daily from P21 to P26, with the last dose occurring 24h prior to collection.

##### Tissue collection and enrichment for biotinylated proteins

For all steps below, low adhesion Eppendorf tubes (USA Scientific 1415-2600) and HPLC grade water (Fisher W71) were used. Cortices were collected at P27 as described previously for fresh tissue, then frozen at -80C. Each sample consisted of two cortices from one mouse. All 16 samples (12 experimental ER-TurboID, 3 mice per genotype; 4 control smFP, 1 mouse per genotype; samples were from the four genotypes WT, WT;Smad4 cKO, FXS, FXS;Smad4 cKO) were then thawed on ice and processed in parallel. Each sample was added to 1mL of NP40 lysis buffer (50mM Tris pH 7.5, 150mM NaCl, 5mM EDTA, 0.5% NP40 [Thermo 11668-019]) with 1:100 protease and phosphatase inhibitors and 1mM DTT in a Precellys homogenization tube (Bertin P000973-LYSK0-A.0), then lysed with a Cryolys Evolution/Precellys Evolution tissue homogenizer (Bertin K002198-PEVO0-A.0; 5800rpm, 10C, 2x30 s, 15 s rest, cryolyse ON) and centrifuged at 20000 g for 15 min at 4C. The supernatant was decanted and protein concentration measured with BCA. 5mg per sample of lysate was added to 50ul Protein G beads (Cytiva 28951379) pre-bound with 10ul anti-PCB (Abcam 110314, rotating in 500ul NP40 lysis buffer at room temperature for 3 hours then washed 1x with 500ul 50mM Tris, 150mM NaCl pH 7.5). For one sample (WT;Smad4cKO1), 3mg lysate was used as input and all reagents were adjusted proportionally. Each tube was leveled to 1mL total volume with NP40 lysis buffer with 1:100 protease and phosphatase inhibitors, then rotated at 4C for 24h. The solution supernatant was separated from Protein G beads with a magnetic stand (DynaMag-2 Magnet, Thermo 12321D), and the supernatant was added to 200ul of washed streptavidin magnetic beads (Thermo 88817). DTT was added to 0.5mM, and the solution was rotated for 1hr at room temperature, then 16hr at 4C. The magnetic beads were isolated with a magnetic stand, then washed at room temperature and eluted. Specifically, samples were washed with 200ul NP40 lysis solution (2 min), transferred to a new tube, washed with 1mL NP40 lysis solution with 1mM DTT (2 min), 1mL 1M KCl (2 min), 1mL 2M urea in 10mM Tris-HCl (10 s), 1mL NP40 lysis solution (2 min, transfer to a new tube), and 1mL NP40 lysis solution. Biotinylated proteins were eluted with 200ul elution buffer (1x PLB, 20mM DTT, 2mM biotin), for 5 min at 95C.

##### Validation western blotting

Western blotting was performed as described above with modifications as below. 15-well gels were used, and both pre-enrichment lysates (10ug/sample) and eluted biotinylated proteins were analyzed. Lysates were blotted for Streptavidin 680 (no primary antibody overnight incubation, 1:3000 in blocking buffer + 0.1% Tween, 1hr room temperature), as further described in the Western blotting section.

##### Immunohistochemistry and analysis: virus validation

The penetrance and specificity of the GfaABC1D-ER-TurboID and GfaABC1D-smFP viruses were quantified. For GfaABC1D-ER-TurboID, virus was injected and biotin administered as described above, for three mice total, with tissue collected at P20 prior to biotin administration, as well as at P27 after six days of biotin. For GfaABC1D-smFP, two mice were injected with virus at P12, treated with biotin from P21-25, and collected at P26; one mouse was injected with virus at P15 and collected at P28. Sagittal brain sections of 16 um were obtained from mice injected with each virus (1.8mm lateral from midline). Slices were blocked and permeabilized in blocking buffer (10% goat serum [Life Tech 16210072] in PBS with 0.3% Triton and 100mM L-lysine) for one hour at room temperature. Primary antibodies were diluted in antibody dilution solution (5% goat serum in PBS with 0.3% Triton and 100mM L-lysine) and stained overnight in a humidified chamber at 4C. Antibodies for the astrocyte marker SOX9 (1:1000, rabbit, Abcam ab185966), HA antibody (1:500, rat, Roche Clone 3F10) were used. The next day slices were washed three times in PBS with 0.2% Triton at 5, 10, and 15 minutes. Secondary antibody labeling was performed in antibody solution (5% goat serum in PBS with 0.3% Triton and 100mM L-lysine) and for two hours at room temperature. Appropriate secondary antibodies were used (1:500, goat anti-rat Alexa 488 Invitrogen A11006; 1:500, goat anti-rabbit Alexa 555 Invitrogen A21429; 1:500 streptavidin Alexa 647 Invitrogen S32357). Washes in PBS with 0.2% Triton were performed three times at 5, 10, and 15 minutes. DAPI was applied (1:1000) for 5 minutes followed by a PBS wash. Coverslips were mounted with DAPI slowfade (Thermo S36939). Images were obtained on an AxioImager.Z2 using a 20x objective (NA = 0.8) as 14-bit images. Tiled (2x2) z-stacks (2637 x 2637 pixels, pixel size 0.323 um x 0.323 um) spanning all layers of visual cortex were acquired with 6 z-steps of 1 um each, totaling a width of 5 um. Three slices per mouse were imaged with three mice total. The images were stitched and maximum intensity projections were created in Zen software. The images were then imported into ImageJ for cell counting analysis using the cell counter plug-in. The number of SOX9+ cells were counted in each image to obtain the total number of astrocytes, respectively. Then the number of HA+ and HA+/SOX9+ cells were counted. The viral penetrance was quantified as the percentage of SOX9+ cells that were also HA+ for each virus. The viral specificity was quantified as the percentage of HA+ cells that were also SOX9+ for each virus. Images shown in **Fig. 4B** and **Supplementary Fig. 8A,B** are maximum intensity projections.

##### Tandem mass tagging mass spectrometry

Samples were precipitated by methanol/chloroform and redissolved in 8 M urea/100 mM TEAB, pH 8.5. Proteins were reduced with 5 mM tris(2-carboxyethyl)phosphine hydrochloride (TCEP, Sigma-Aldrich 68957) and alkylated with 10 mM chloroacetamide (Sigma-Aldrich 22790). Proteins were digested overnight at 37C in 2 M urea/100 mM TEAB, pH 8.5, with trypsin (Promega V5280). The digested peptides were labeled with 16-plex TMT (Thermo A44522, lot YB368939), pooled samples were fractionated by basic reversed phase (Thermo 84868). Each of the 16 channels in the 16-plex was used to label a separate sample. The 16 samples consisted of 12 experimental samples (3 mice per genotype) injected with virus expressing ER-TurboID, and 4 control samples (1 mouse per genotype) injected with virus expressing control protein smFP. Samples were from the four genotypes WT, WT;Smad4 cKO, FXS, and FXS;Smad4 cKO.

The TMT labeled samples were analyzed on a Orbitrap Eclipse Tribrid mass spectrometer (Thermo). Samples were injected directly onto a 25 cm, 100 μm ID column packed with BEH 1.7

μm C18 resin (Waters SKU186002352). Samples were separated at a flow rate of 300 nL/min on an EasynLC 1200 (Thermo LC140). Buffer A and B were 0.1% formic acid in water and 90% acetonitrile, respectively. A gradient of 1–15% B over 30 min, an increase to 45% B over 120 min, an increase to 100% B over 20 min and held at 100% B for 10 min was used for a 180 min total run time.

Peptides were eluted directly from the tip of the column and nanosprayed directly into the mass spectrometer by application of 2.5 kV voltage at the back of the column. The Eclipse was operated in a data dependent mode. Full MS1 scans were collected in the Orbitrap at 120k resolution. The cycle time was set to 3 s, and within this 3 s the most abundant ions per scan were selected for CID MS/MS in the ion trap. The TMT samples were analyzed by MS3 analysis, and multinotch isolation (SPS3) was utilized for detection of TMT reporter ions at 60k resolution.^111^ Monoisotopic precursor selection was enabled and dynamic exclusion was used with exclusion duration of 60s. Protein and peptide identification were done with Integrated Proteomics Pipeline – IP2 (Integrated Proteomics Applications, v6.0.5). Tandem mass spectra were extracted from raw files using RawConverter^112^ and searched with ProLuCID^113^ against Uniprot mouse database. The search space included all fully-tryptic and half-tryptic peptide candidates. Carbamidomethylation on cysteine and TMT on lysine and peptide N-term were considered as static modifications. Data were searched with 50 ppm precursor ion tolerance and 600 ppm fragment ion tolerance. Identified proteins were filtered using DTASelect^114^ utilizing a target-decoy database search strategy to control the false discovery rate to 1% at the protein level.^115^ Quantitative analysis of TMT was done with Census,^116^ filtering reporter ions with 10 ppm mass tolerance and 0.6 isobaric purity filter.

##### Proteomic analysis

Proteins were filtered for contaminants and duplicates as follows: proteins with identical gene names were removed, proteins annotated as “contaminant” were removed, and streptavidin was removed. For the one sample (WT;Smad4cKO1) for which a different amount of input was used, the TMT signal was normalized for the input amount (multiplication by a factor of 1.71, after accounting for aliquots used for QC testing). Proteomic data was analyzed with differentially expressed protein criteria: p-value < 0.05 by unpaired t-test and |fold change (FC)| > 1.25. Heatmaps were created using R (v4.2.2) via RStudio (2022.12.0) using the Pheatmap package and Graphpad Prism (v8.0). Comparison to FMRP targets used reference database in Darnell et al., 2011.^52^ Volcano plots were created with package ggplot2 (v3.4.0).^117^

Gene set enrichment analysis (GSEA) was performed to detect which pathways and gene sets were enriched in Fmr1 KO and Smad4 cKO using the package ClusterProfiler (v4.6.0).^101^ The following gene sets from mouse MSigDB^102^ were used: hallmark pathways, canonical pathways (including Reactome,^103^ KEGG,^104^ Wikipathways,^105^ and BioCarta^106^), and gene ontology.^107,108^ Proteins with Turbo/smFP ratio > 1.75 were used for GSEA, with ranking based on sign(L2FC)*-log10(p-value), for each comparison. For GSEA pathway analysis only, peptides mapped to multiple protein isoforms were only counted once as the alphanumerically first protein isoform, in order to avoid overweighting single peptides. Highlighted pathways in the visualization are the top three down- and upregulated (lowest and highest normalized enrichment score [NES]) hallmark pathways in each comparison (**Fig. 6D**). Cellular compartment of differentially expressed proteins was determined via UniProt annotations. Rank-rank hypergeometric plot was generated using the package RRHO v1.38.0.^118^

The ER-TurboID proteome was visualized with Cytoscape v3.10.1,^119^ with nodes labeled as the gene names for the top 200 enriched proteins (Turbo/smFP ratio). Node size for detected proteins was set as proportional to average log10(TMT intensity) across all 12 experimental samples. Clustergrams were created via Uniprot, STRING, and GeneCards database annotations. For neurologic disease association, only proteins classified as “causative mutation” in GeneCards were included.^120^ Astrocyte enrichment was determined referencing the external database in Zhang et al., 2014.^53^ Cellular compartment gene ontology^107,108^ on the ER-TurboID proteome was conducted via overrepresentation analysis with the package ClusterProfiler (v4.6.0).^101^

For testing for overlap of known FMRP targets with up- and downregulated proteins in Fmr1 KO (Comparison 1), the list of 842 genes from Darnell et al. 2011^52^ was used, and statistics were done with Fisher’s exact test with genome size 24,252 protein-coding *M. Musculus* genes.

##### Immunohistochemical synaptic staining, imaging and analysis

Littermate WT and WT;Smad4 cKO pairs and littermate FXS and FXS;Smad4 cKO pairs at P28 were used for experiments, and 8 independent experiments were performed, each with one mouse of each genotype (biological replicates) processed in parallel and blind to genotype from sectioning onwards. Coronal brain sections of PFA-fixed tissue of 18 um thickness were obtained using a cryostat (Hacker Industries OTF5000) (2.7mm posterior to bregma). Slices were blocked and permeabilized in blocking buffer (10% goat serum in PBS with 0.3% Triton and 100mM L-lysine) for one hour at room temperature. Primary antibodies were diluted in antibody dilution solution (5% goat serum in PBS with 0.3% Triton and 100mM L-lysine) and stained overnight in a humidified chamber at 4C. The primary antibodies for inhibitory synaptic staining were Vgat (1:250, guinea pig, Synaptic Systems 131004) and gephyrin (1:500, rabbit, Synaptic Systems 147008), and for excitatory synaptic staining were Vglut1 (1:2000, guinea pig, Millipore Sigma AB5905) and PSD95 (1:200, rabbit, Thermo 51-6900). The next day the slides were washed in PBS with 0.2% Triton 3 times, for 5 minutes each. Appropriate secondary antibodies were used in antibody dilution solution (1:500, goat anti-guinea pig Alexa 594, Invitrogen A11076; 1:500, goat anti-rabbit Alexa 488, Invitrogen A11034). Secondary antibodies were applied for two hours at room temperature followed by three washes at 5, 10, and 15 minutes in PBS with 0.2% Triton. DAPI was applied (1:1000) for five minutes followed by a PBS wash. Coverslips were mounted with SlowFade Gold Antifade Mountant with DAPI (Invitrogen, S36939). As negative controls, the primary antibodies were omitted and the brain slices were incubated with just secondary antibodies. Images were obtained using a 63x objective (NA = 1.4) with 1.8x zoom on an LSM 880 confocal microscope with Airyscan module using super-resolution mode at 1764 x 1764 pixels, 0.04 um x 0.04 um pixel size, with pixel dwell time 2.38us, as an 8-bit image (16-bit after Airyscan processing). The laser settings and gain were maintained constant for all experiments. Z-stacks of 0.18 um increments were obtained in Layer 2/3 with a total of 20 z-steps per image, from a depth of 2.7 um to 6.3 um from tissue surface. Within each experiment, each mouse was imaged and analyzed on 2-5 independent brain sections (technical replicates), that were then averaged per mouse. Quantification of colocalized puncta was performed as previously described using the Imaris software (Bitplane) spots tool and colocalization algorithm^10,31,42^. Spot sizes were set to 0.35 um for Vgat and Vglut1 puncta and 0.3 um for Gephyrin and PSD95 puncta with a center-to-center colocalization distance of 0.6 um. Thus spots from the two channels that were within 0.6 um from each other’s center were quantified as a synapse. Quality threshold for puncta detection was maintained constant within each experiment. Images shown in **Fig. 7A,C** are maximum orthogonal projections of the 20 z-steps.

##### Quantification and statistical analysis

Statistical analysis and data visualization were performed in GraphPad Prism 8 and R (v4.2.2). All tests were two-tailed. Data was tested for normality using either Shapiro-Wilk or Kolmogorov-Smirnov tests. For normally distributed data, unless otherwise noted, unpaired t-tests were used to compare two groups and 2-way ANOVAs were used to compare multiple groups and treatments, with post-hoc comparison testing pre-determined by the experimental design: Sidak’s multiple comparison test was used to assess genotype changes across multiple cortical layers (**Supplementary Fig. 1A-D**, **2D,E**, **3B-D,F-H**), while Tukey’s multiple comparison test was used in all other instances to compare each group to each other group. Comparisons of categorical data were performed with Chi-Squared test (more than two categories) and Fisher’s Exact test (two categories). Least-squares linear regression was used to calculate Pearson’s correlation coefficients. Comparisons of L2FC distributions of large gene sets were performed with z-tests. All bar graphs and line graphs represent mean ± standard error of the mean (SEM), with individual data points representing mice unless otherwise indicated. Figures were generated using Adobe Illustrator 2022 and BioRender. Exact p values reported on graphs.

## Materials availability

The GfaABC1D-smFP-HA, GfaABC1D-ER-TurboID-HA, GfaABC1D-sec61b-TurboID-HA, GfaABC1D-cyto-TurboID-HA, and GfaABC1D-C1(1-29)-ER-TurboID-HA pAAV constructs were generated for this study. Original plasmids are available upon request.

## Data availability

The data that support the findings of this study are available from the lead contact upon reasonable request. Deposited RNA sequencing and proteomics data will be made available upon publication.

## Acknowledgements

We thank Joseph Hash for technical assistance with the mouse colony. We thank J. Gleeson, S. Ackerman, Y. Zou, R. Daneman, L. Sancho, the Allen lab, and members of Salk MNL for discussions. Biorender was used to create cartoon schematics. JD is supported by a FRAXA Research Foundation Fellowship and F30 HD106699 from the National Institute of Child Health and Human Development. NJA is supported by the Chan Zuckerberg Initiative. SBK was supported by a URS Eureka! Research Scholarship. LLB is supported by a Hewitt-Eckhart postdoctoral fellowship, and ANB is supported by NIA 1K99AG081536-01. This work was supported by core facilities of the Salk Institute supported by NIH-NCI CCSG: P30 CA01495 and NIH-NIA San Diego Nathan Shock Center P30 AG068635, including the Mass Spectrometry Core, also supported by the Helmsley Center for Genomic Medicine; the Integrative Genomics and Bioinformatics Core, also supported by the Helmsley Trust; In Vivo Scientific Services; the GT3 core, also supported by NINDS Core Grant R24NS092943 and the NEI; Flow Cytometry, also supported by SIG S10-OD023689 (Aria Fusion cell sorter); and the Waitt Advanced Biophotonics Core, also supported by the Waitt Foundation. This publication includes data generated at the UC San Diego IGM Genomics Center using an Illumina NovaSeq 6000 purchased with funding from the NIH SIG grant #S10 OD026929. The funders had no role in study design, data collection and analysis, decision to publish, or preparation of the manuscript.

## Author Contributions

JDD and NJA designed experiments, analyzed data, and wrote the manuscript with input from all authors. NJA conceived the project. LLB and ANB provided technical guidance on transcriptomics and proteomics respectively. SBK performed experiments and analyzed data. AFMP and JKD performed MS experiments.

## Materials & Correspondence

Correspondence and request for materials should be addressed to Nicola J. Allen.

## Supplementary Data Tables

**Supplemental Data Table 1.** Data from RNA sequencing experiments. Displaying TPM values for each gene for each sample.

**Supplemental Data Table 2.** Differentially expressed genes (DEGs) from RNA sequencing, based on criteria: average TPM > 1 in WT samples, Benjamini-Hochberg (BH) adjusted p-value < 0.05, |fold change (FC)| > 1.5.

**Supplemental Data Table 3.** Gene set enrichment analysis (GSEA) data on RNA sequencing. Enrichment of gene sets and pathways from MSigDB, Gene Ontology (GO), Reactome, KEGG, Wikipathway, and BioCarta. Genes with TPM > 1 in every sample were used for GSEA, with ranking based on sign(log2 fold change)*-log10(p-value), for each comparison.

**Supplemental Data Table 4.** Gene lists and associated RNAseq z-score averaged by genotype, for the glycolysis/gluconeogenesis set (KEGG M11521) and the oxidative phosphorylation set (MSigDB MM3893).

**Supplemental Data Table 5.** Gene list and pairwise comparisons in RNA sequencing data for BMP pathway components.

**Supplemental Data Table 6.** Data from proteomics experiment. Displaying TMT raw signal values for each protein for each sample, average TMT signal per genotype, and enrichment of experimental over control samples.

**Supplemental Data Table 7.** Differentially expressed proteins (DEPs) from proteomics, based on criteria: p-value < 0.05 using t-test, |fold change (FC)| > 1.25.

**Supplemental Data Table 8.** Gene set enrichment analysis (GSEA) data on proteomics. Enrichment of sets and pathways from MSigDB, Gene Ontology (GO), Reactome, KEGG, Wikipathway, and BioCarta. Proteins with Turbo/smFP enrichment ratio > 1.75 were used for GSEA, with ranking based on sign(log2 fold change)*-log10(p-value), for each comparison.

**Supplemental Data Table 9.** Statistical tests and summary statistics from studies.

---

**Supplementary Figure 1.**
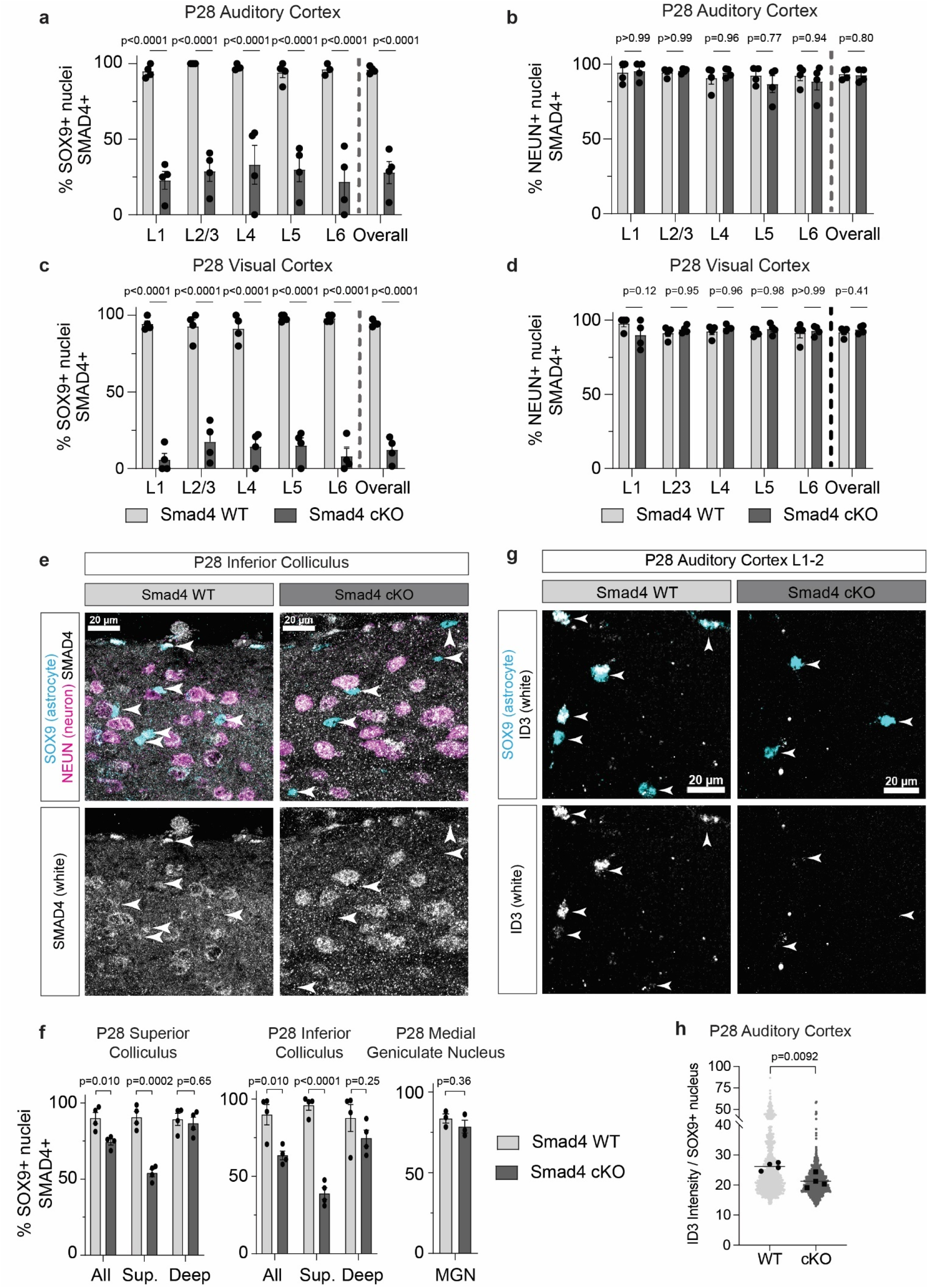
Additional immunohistochemical validation and characterization of astrocyte Smad4 cKO. **a**. Smad4 cKO effectiveness in P28 auditory cortex. N = 4 mice per genotype. Statistics for layers by 2-way ANOVA with Sidak’s test for multiple comparisons. Statistics for overall by t-test; data for overall is the same as in Fig. 1d. **b**. Smad4 cKO has no effect on neuron SMAD4 immunopositivity in P28 auditory cortex. N = 4 mice per genotype. Statistics for layers by 2-way ANOVA with Sidak’s test for multiple comparisons. Statistics for overall by t-test. **c**. Smad4 cKO effectiveness in P28 visual cortex. N = 4 mice per genotype. Statistics for layers by 2-way ANOVA with Sidak’s test for multiple comparisons. Statistics for overall by t-test. **d**. Smad4 cKO has no effect on neuron SMAD4 immunopositivity in P28 visual cortex. N = 4 mice per genotype. Statistics for layers by 2-way ANOVA with Sidak’s test for multiple comparisons. Statistics for overall by t-test. **e**. Smad4 cKO achieves KO in P28 inferior colliculus. Immunohistochemistry for astrocyte nuclear marker SOX9, neuronal nuclear marker NEUN, and SMAD4 shows removal of SMAD4 specifically in some astrocytes. White arrowheads indicate locations of SOX9+ astrocyte nuclei. **f**. Quantification of Smad4 cKO at P28 in superior colliculus, inferior colliculus, and medial geniculate nucleus of the thalamus. For the colliculi, SOX9+ cells were quantified both separated by superficial (“sup”) vs. deep, as well as altogether (“all”). Superficially located SOX9+ cells were defined as located within 100um of the dorsal pial surface, and deep SOX9+ cells were defined as located greater than 100um from the dorsal pial surface. N = 3-4 mice per genotype per age. Statistics by t-test. **g**. Smad4 cKO downregulates ID3 in astrocytes. Example images of L1-2 P28 auditory cortex, immunohistochemistry for ID3 and SOX9 (astrocyte nuclei). White arrowheads indicate locations of SOX9+ astrocyte nuclei. **h**. Quantification of ID3 intensity within SOX9 nuclei across P28 auditory cortex. Black dots denote average for one mouse, grey dots denote individual SOX9+ nuclei. N = 4 independent experiments, each quantifying one littermate pair. Statistics by t-test on mice.

**Supplementary Figure 2.**
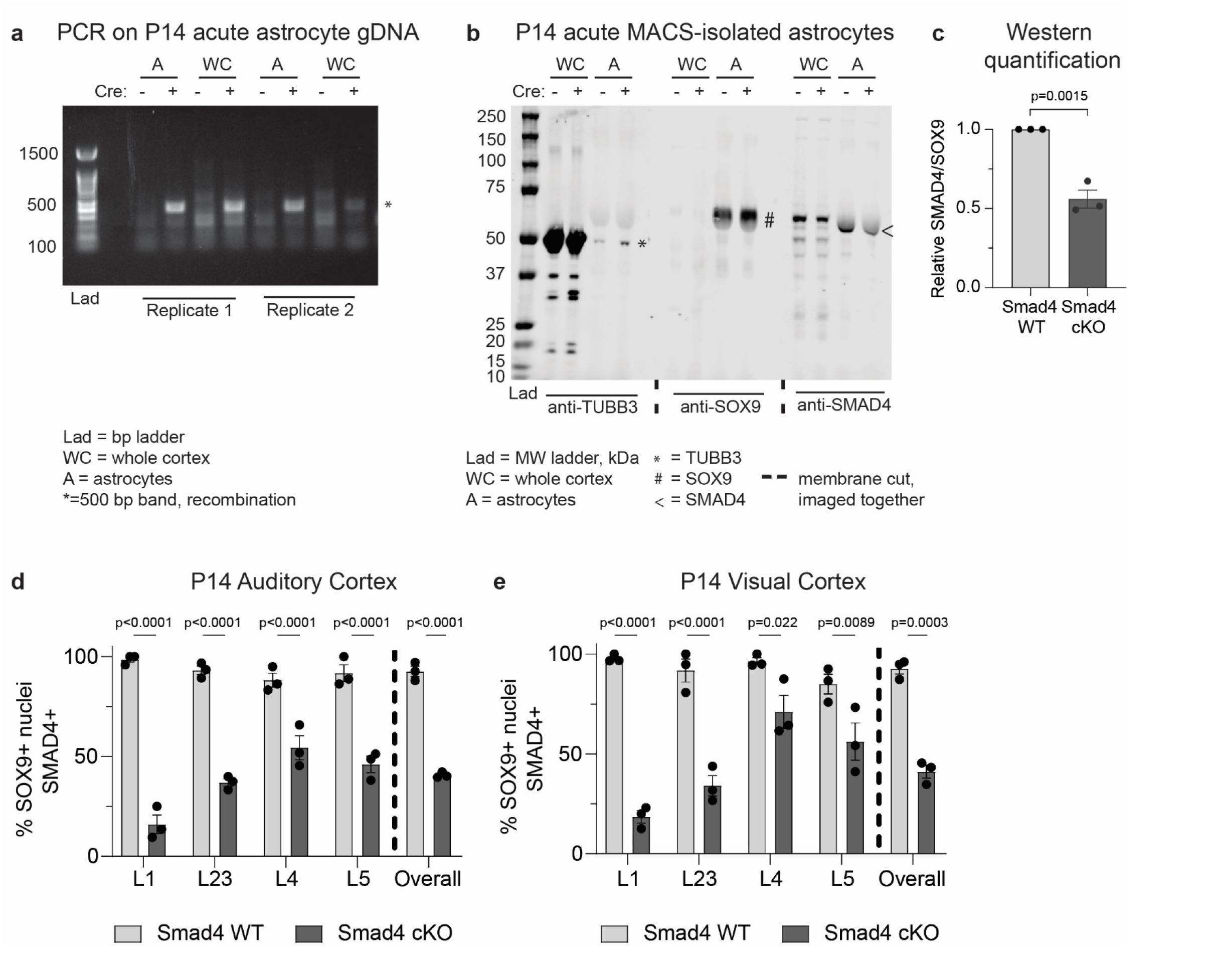
Additional validation and characterization of astrocyte Smad4 cKO at P14. **a**. PCR for recombined Smad4 sequence (see primer in methods) in genomic DNA isolated from whole cortex (WC) and magnetic-activated cell sorting (MACS)-purified acute whole cortical astrocytes with and without Smad4 cKO at P14. N = 2 independent experiments. **b**. Whole cortex (WC) and MACS-purified acute whole cortical astrocytes (A) with and without Smad4 cKO isolated at P14 and probed by western blot for neuron marker TUBB3, astrocyte marker SOX9, and SMAD4. Cortices from 2-3 mice were combined in one MACS experiment for each sample. **c**. Quantification of **b**. N = 3 independent experiments. Statistics by t-test. **d**. Smad4 cKO effectiveness in P14 auditory cortex. N = 3 mice per genotype. Statistics for layers by 2-way ANOVA with Sidak’s test for multiple comparisons. Statistics for overall by t-test. **e**. Smad4 cKO effectiveness in P14 visual cortex. N = 3 mice per genotype. Statistics for layers by 2-way ANOVA with Sidak’s test for multiple comparisons. Statistics for overall by t-test.

**Supplementary Figure 3.**
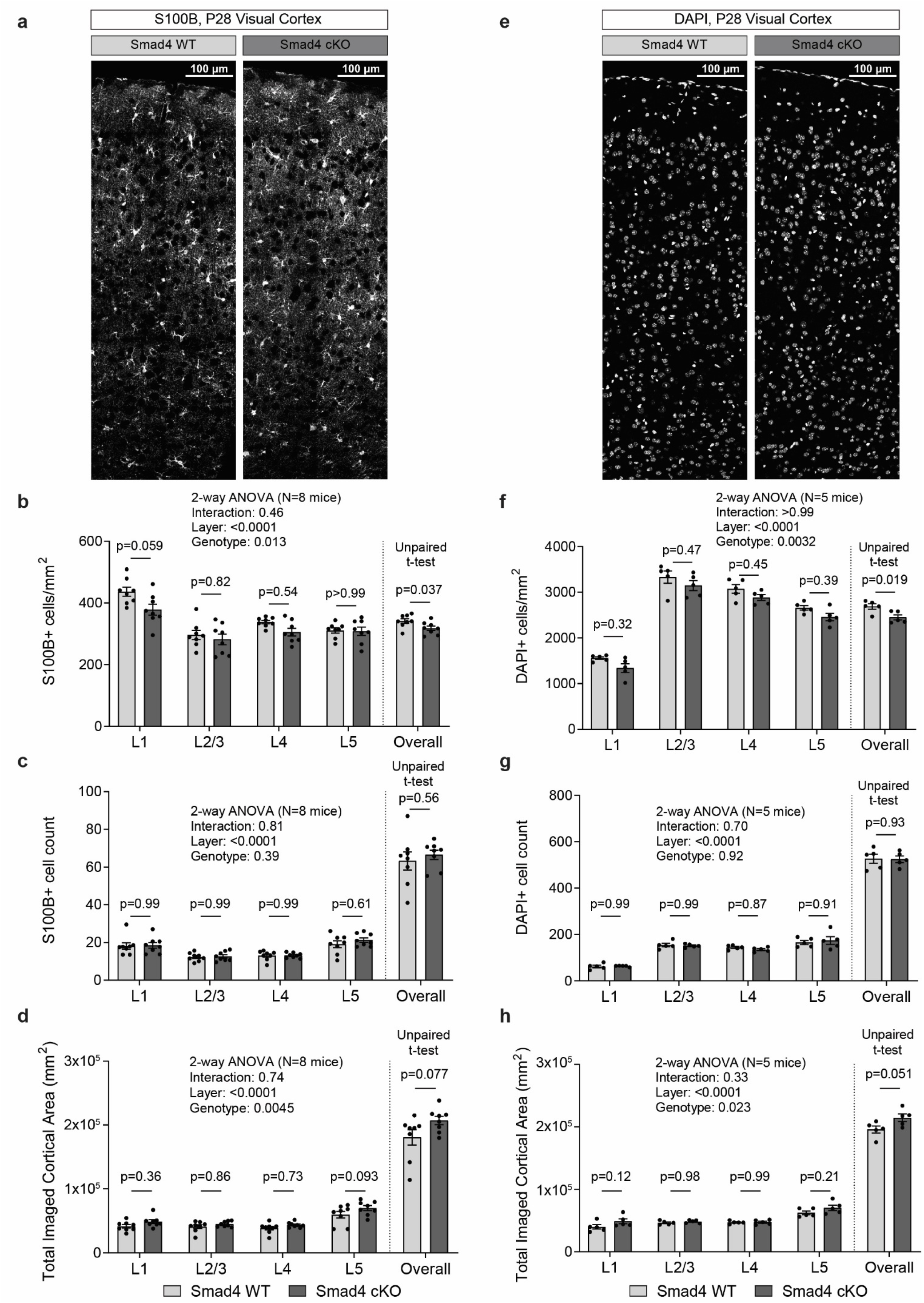
Characterization of astrocyte Smad4 cKO cell density and cortical area. **a**. Immunohistochemistry for astrocyte cytosolic marker S100B in P28 visual cortex. **b-d**. Astrocyte marker S100B density **b**., absolute S100B+ cell count **c**., and imaged cortical area **d**. by layer. N = 8 mice pairs. Statistics for cortical layers by 2-way ANOVA with Sidak’s test for multiple comparisons, statistics for overall by t-test. **e**. Nuclear marker DAPI in P28 visual cortex. **f-h**. Nuclear marker DAPI density **f**., absolute DAPI+ nuclei count **g**., and imaged cortical area **h**. by layer. N = 5 mice pairs. Statistics for cortical layers by 2-way ANOVA with Sidak’s test for multiple comparisons, statistics for overall by t-test.

**Supplementary Figure 4.**
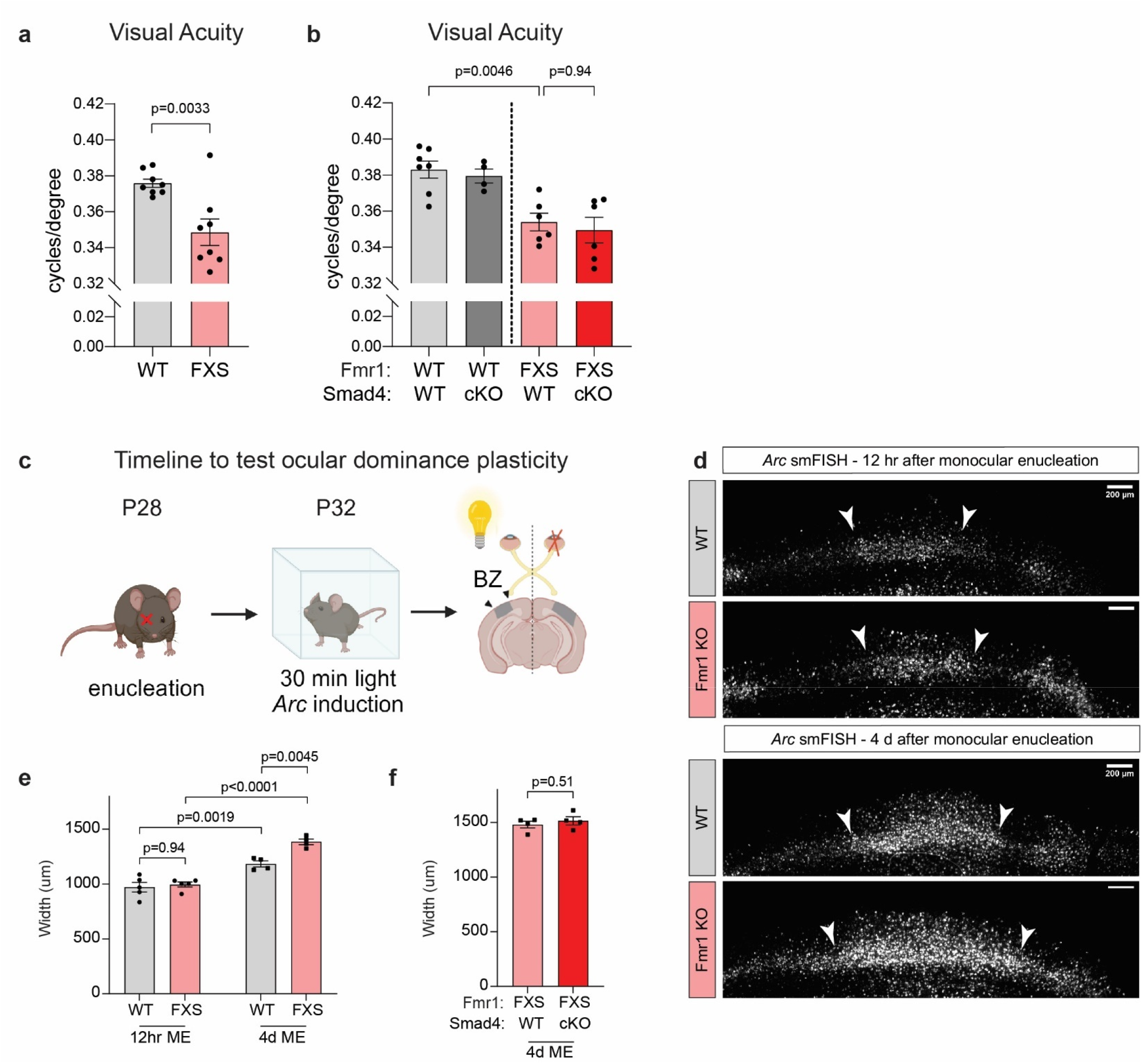
Smad4 cKO does not improve Fmr1 KO visual acuity or ocular dominance plasticity deficits. **a**. Visual acuity in the Fmr1 KO line. N = 8 Fmr1 WT, 8 Fmr1 KO. Statistics by t-test. **b**. Visual acuity in the Fmr1 KO;Smad4 cKO line. N = 7 WT, 4 WT;Smad4 cKO, 6 FXS, 6 FXS;Smad4 cKO. Statistics by 2-way ANOVA with Tukey’s test for multiple comparisons. **c**. Timeline for Arc assay. **d**. Example images of *Arc* smFISH in binocular zone (BZ) of visual cortex from experiments on the Fmr1 KO line. White arrowheads denote edges of the BZ, used to assess BZ width. **e**. Quantification **d**., N = 5 WT 12hr, 5 Fmr1 KO 12hr, 4 WT 4d, 4 Fmr1 KO 4d. Statistics by 2-way ANOVA with Tukey’s test for multiple comparisons. **f**. Quantification of experiments in the Fmr1 KO;Smad4 cKO line. N = 4 FXS, 4 FXS;Smad4 cKO. Statistics by t-test.

**Supplementary Figure 5.**
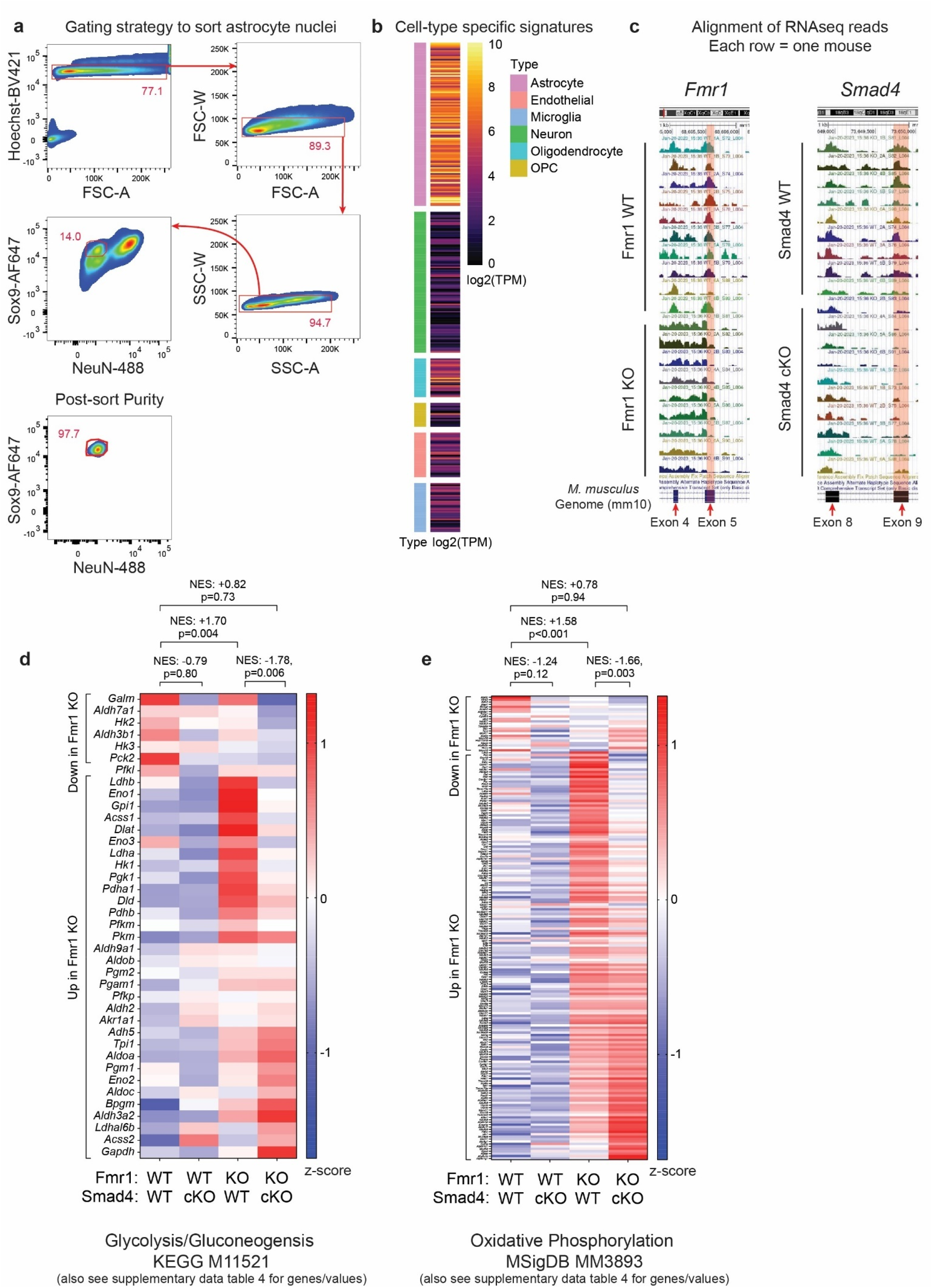
Smad4 cKO moderates Fmr1 KO astrocyte hypermetabolic transcriptional state. **a**. Above, FACS gating strategy to obtain SOX9+NEUN-nuclei for sequencing. Below, example purity plot. **b**. Heatmap of average abundance in nRNAseq for known cell type-specific genes. OPC = oligodendrocyte precursor cell. **c**. Bigwig plots aligning RNAseq reads to the *M. Musculus* genome (mm10). Each sample (one mouse) is one row. Fmr1 KO samples have depletion of reads to the latter half of *Fmr1* exon 5, corresponding to the locus where a neomycin resistance cassette was inserted to create the KO. Smad4 cKO samples have depletion of reads to *Smad4* exon 9, the locus where loxP sites were inserted to create the floxed mouse. The red rectangles denote these loci. **d**. Z-scores of all genes in the “glycolysis/gluconeogenesis” gene set (KEGG M11521), averaged within each genotype, displaying all 39 genes in the set with average TPM > 1 for each genotype group. **e**. Z-scores of all genes in the “oxidative phosphorylation” gene set (MSigDB 3893), averaged within each genotype, displaying all 193 genes in the set with average TPM > 1 for each genotype group.

**Supplementary Figure 6.**
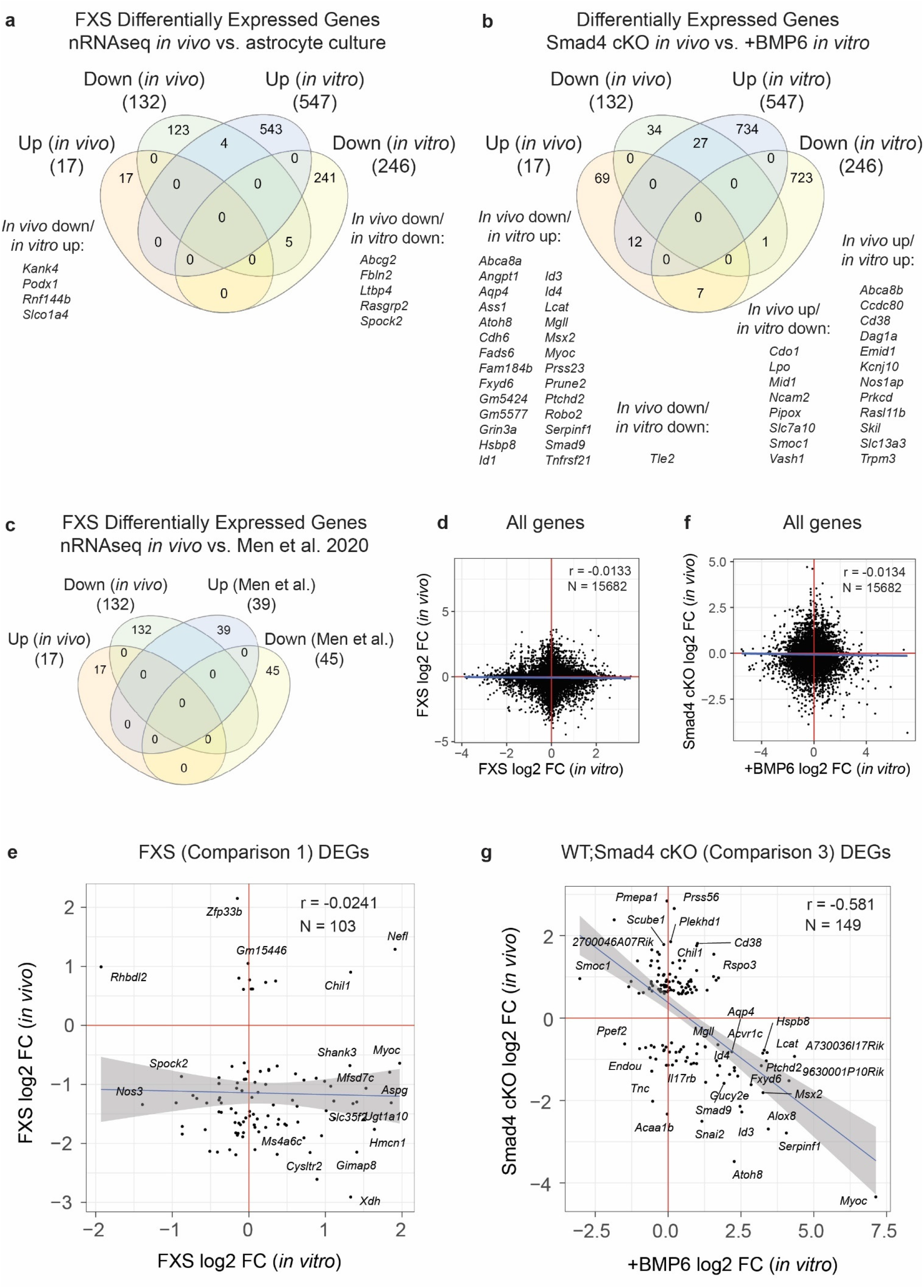
Comparison of *in vivo* transcriptomics with other datasets. **a**. Overlap of Fmr1 KO up- and downregulated genes from *in vivo* RNAseq with up- and downregulated genes from RNAseq of cultured P7 Fmr1 KO astrocytes.^12^ **b**. Overlap of Smad4 cKO up- and downregulated genes from *in vivo* RNAseq with up- and downregulated genes from RNAseq of cultured P7 astrocytes incubated with BMP6.^12^ **c**. Overlap of Fmr1 KO up- and downregulated genes from *in vivo* RNAseq with translating RNAseq profiling of P40 Fmr1 KO astrocytes in the FVB genetic background.^11^ **d**. Scatterplot of L2FCs for all 15,682 genes detected, between *in vivo* RNAseq and cultured Fmr1 KO astrocytes.^12^ **e**. Scatterplot as in **d**., limited to the set of 103 *in vivo* RNAseq Fmr1 KO DEGs also detected *in vitro*.^12^ **f**. Scatterplot of L2FCs for the set of all 18,300 genes between *in vivo* RNAseq Smad4cKO and cultured WT astrocytes incubated with BMP6.^12^ **g**. Scatterplot as in **f**., limited to the set of 149 *in vivo* RNAseq Smad4cKO DEGs also detected *in vitro*.^12^

**Supplementary Figure 7.**
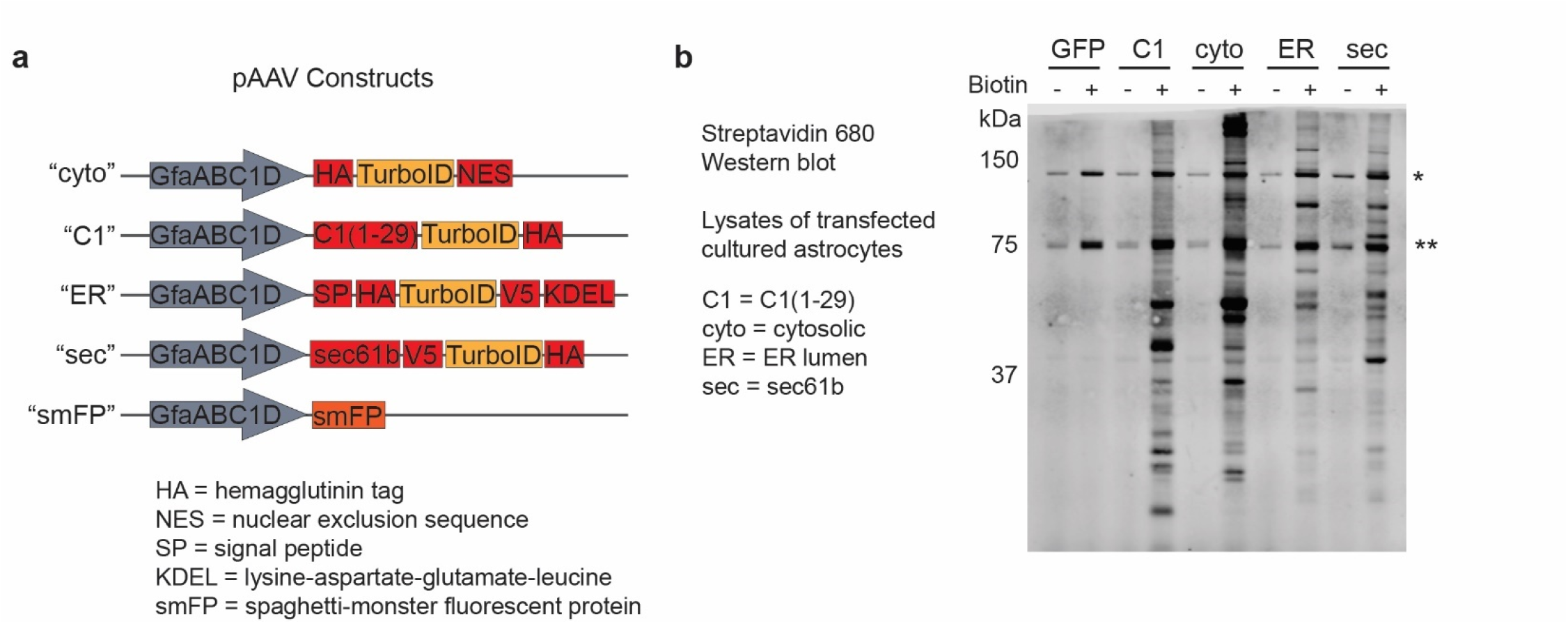
Development of ER-TurboID to study astrocyte secreted and membrane proteins. **a**. Schematics of constructs used: GfaABC1D-HA-TurboID-NES (cytosolic control), GfaABC1D-C1(1-29)-TurboID (ERM facing cytosol), GfaABC1D-HA-TurboID-V5-KDEL (ER lumen), GfaABC1D-sec61b-V5-TurboID-HA (ERM facing lumen), and GfaABC1D-smFP (control protein). **b**. Streptavidin 680 signal on western blot of lysates of cultured astrocytes transfected with the five constructs from **a**. TurboID functionally incorporates biotin nonspecifically when biotin is added to culture media. * = pyruvate carboxylase, known endogenous biotinylated protein; ** = proprionyl-CoA carboxylase alpha chain, known endogenous biotinylated protein.

**Supplementary Figure 8.**
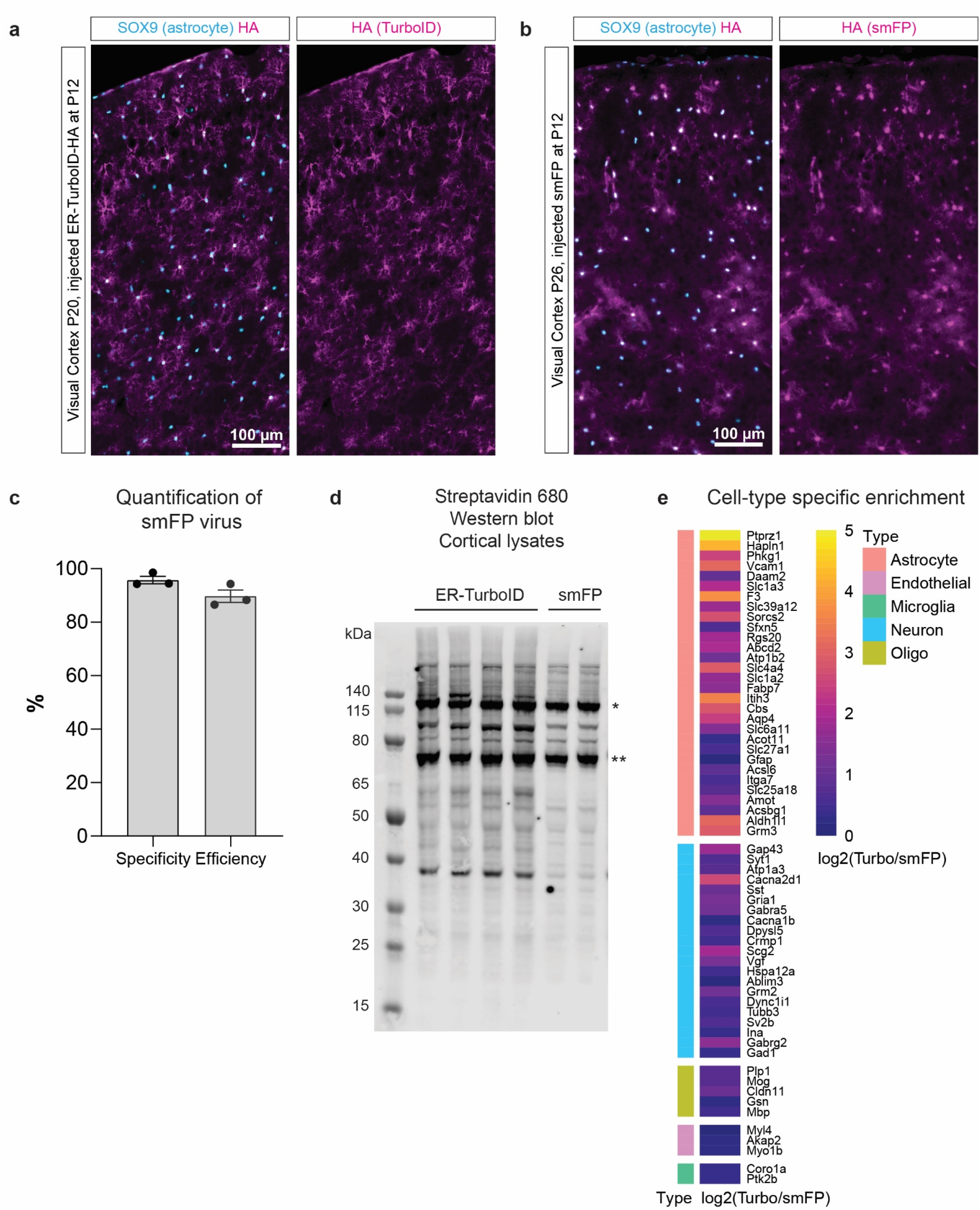
Validation of ER-TurboID for studying astrocyte proteins *in vivo*. **a**. AAV-PHP.eB-packaged ER-TurboID (HA tagged) is highly expressed in astrocytes by P20 after P12 retroorbital injection. Immunohistochemistry in visual cortex for astrocyte nuclear marker SOX9 and HA. **b**. AAV-PHP.eB-packaged smFP is expressed specifically in astrocytes by P26 after P12 retroorbital injection. Immunohistochemistry in visual cortex for astrocyte nuclear marker SOX9 and HA. **c**. Quantification of **b**. Specificity quantified as (# SOX9+HA+ cells)/(# HA+ cells), efficiency quantified as (# SOX9+HA+ cells)/(# SOX9+ cells) for N = 3 mice. **d**. ER-TurboID biotinylates cortical proteins. Streptavidin 680 signal on cortical lysates collected at P27 from mice injected at P12 with ER-TurboID or control protein (smFP) and receiving subcutaneous biotin from P21-26. * = pyruvate carboxylase, known endogenous biotinylated protein; ** = proprionyl-CoA carboxylase alpha chain, known endogenous biotinylated protein. **e**. ER-TurboID TMT signal is enriched for astrocyte proteins over other cell type proteins. Heatmap represents enrichment as log2 of average TurboID TMT signal divided by average control TMT signal.

**Supplementary Figure 9.**
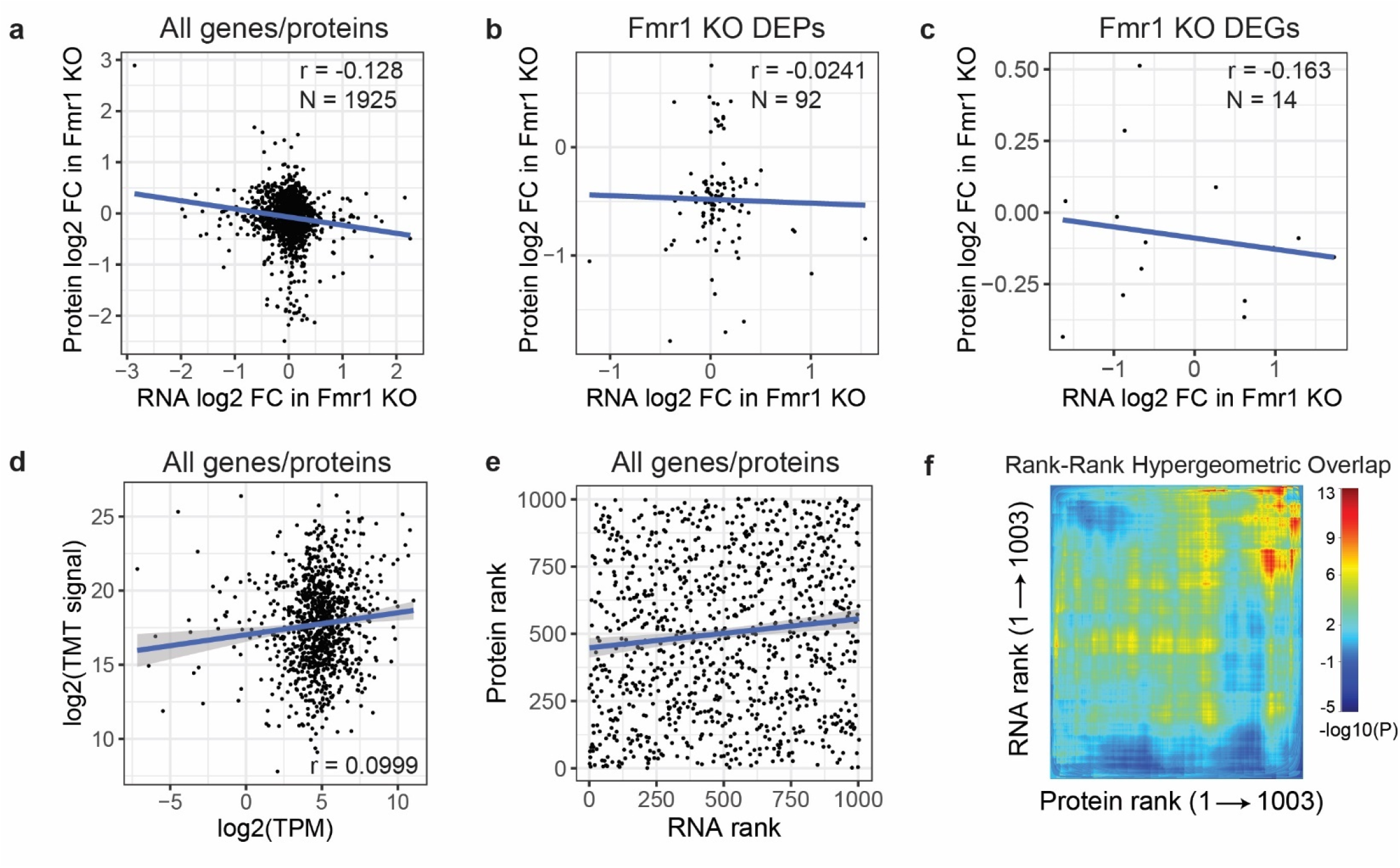
Minimal to slight correlations between *in vivo* transcriptomics and ER-TurboID proteomics. **a**. Plotting log2 FC in the Fmr1 KO comparison, in protein (ER-TurboID) vs. RNA, for all 1925 genes assayed by both methods. **b**. Plotting log2 FC in protein (ER-TurboID) vs. RNA, for all 92 proteomics Fmr1 KO DEPs (p < 0.05, |FC| > 1.25) assayed by both methods. **c**. Plotting log2 FC in protein (ER-TurboID) vs. RNA, for all 14 RNAseq Fmr1 KO DEGs (adj-p < 0.05, |FC| > 1.5, TPM > 1 in WT) assayed by both methods. **d**. Plotting proteomics absolute TMT signal vs. log2 of TPM from RNAseq, for all 1,003 genes assayed by both methods, filtering proteins for only those with average Turbo/Control TMT ratio of over 1.5. **e**. Plotting protein prevalence rank by absolute TMT signal vs. RNA rank by TPM, for all 1003 genes assayed by both methods, filtering proteins for only those with average Turbo/Control TMT ratio of over 1.5. Pearson’s r = 0.108. **f**. Rank-rank hypergeometric overlap map using data from **e**.

**Supplementary Figure 10.**
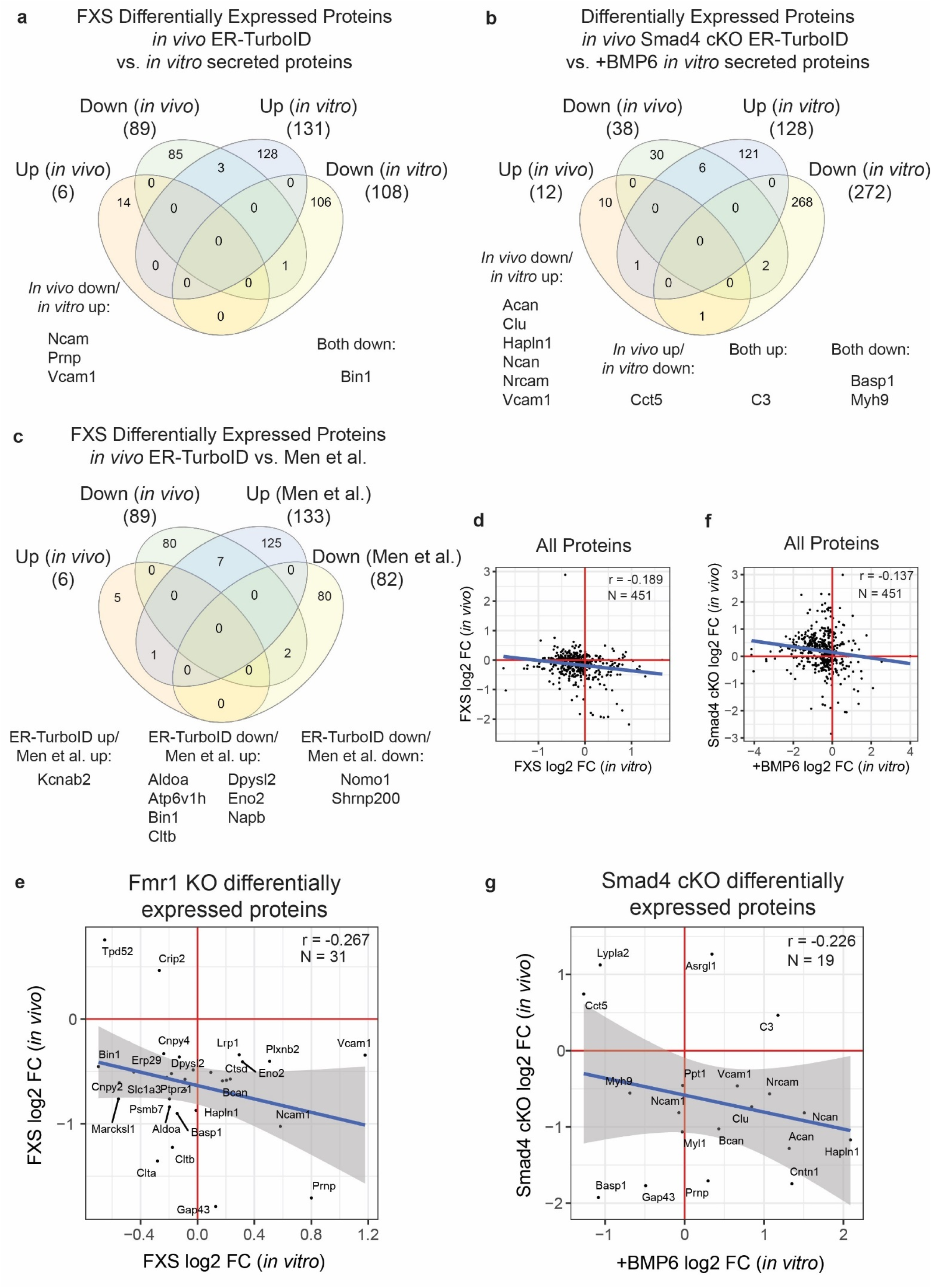
Comparison of ER-TurboID proteomics with other datasets. **a**. Overlap of Fmr1 KO up- and downregulated proteins from ER-TurboID proteomics with up- and downregulated proteins from conditioned media of cultured Fmr1 KO astrocytes.^12^ **b**. Overlap of Smad4 cKO up- and downregulated proteins from ER-TurboID proteomics with up- and downregulated proteins from conditioned media of cultured WT astrocytes incubated with BMP6.^12^ **c**. Overlap of Fmr1 KO up- and downregulated proteins from ER-TurboID proteomics with up- and downregulated proteins from FACS-sorted P40 Fmr1 KO astrocytes in the FVB genetic background.^11^ **d**. Correlations of Fmr1 KO L2FCs for all 451 proteins detected in both ER-TurboID proteomics and cultured Fmr1 KO astrocytes.^12^ **e**. Correlations as in **d**., limited to the set of 31 ER-TurboID proteomics Fmr1 KO DEPs detected *in vitro*.^12^ **f**. Correlations for all 451 proteins detected in both datasets, between L2FCs in Smad4 cKO in ER-TurboID proteomics vs. WT astrocytes incubated with BMP6.^12^ **g**. Correlations as in **f**., limited to the set of 19 ER-TurboID proteomics Smad4 cKO DEPs detected *in vitro*.^12^

**Supplementary Figure 11.**
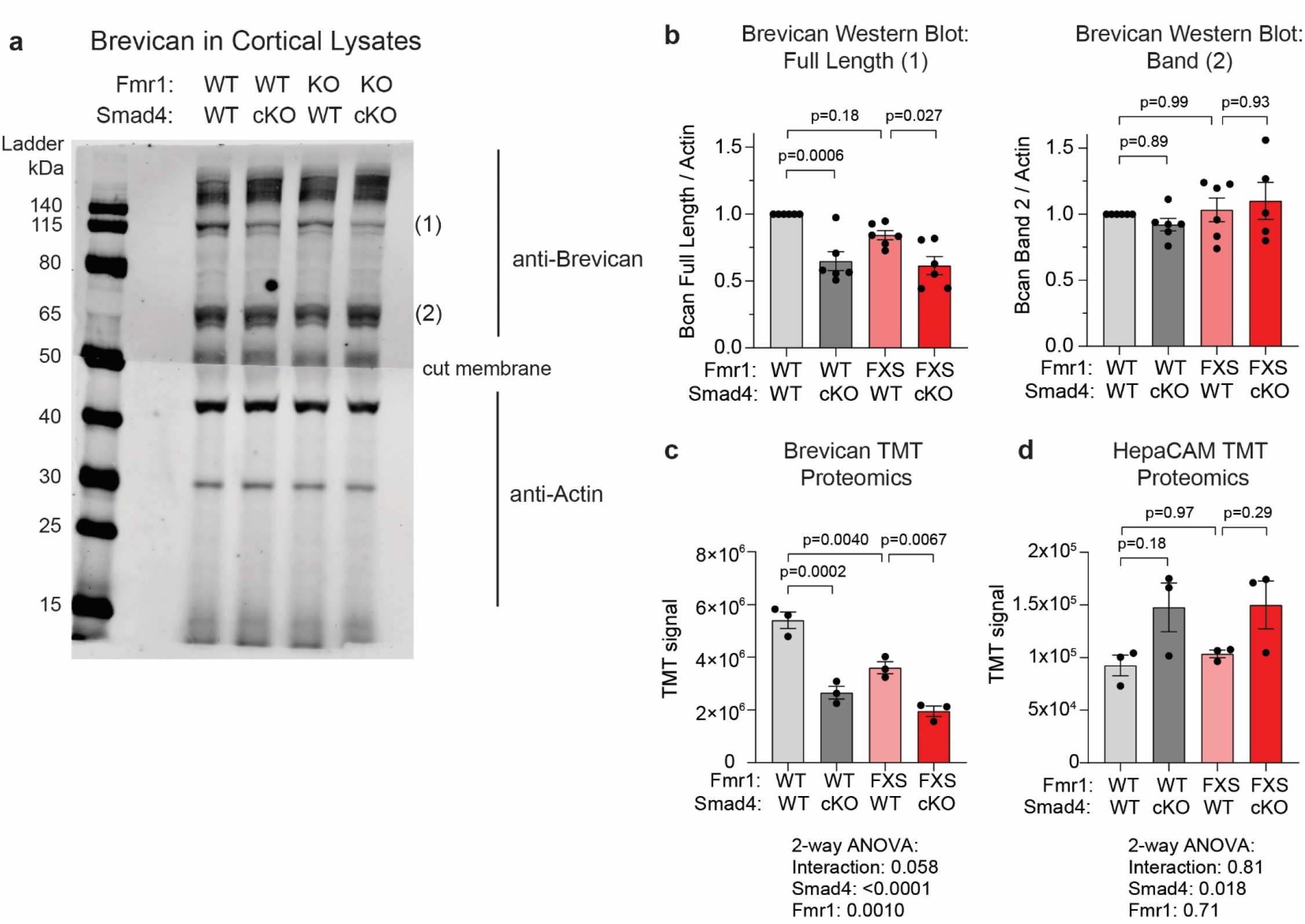
Proteomic TMT data and western blot validation of ER-TurboID-identified differentially expressed proteins brevican and hepaCAM. **a**. Western blotting on cortical lysates for brevican and actin. **b**. Quantification of two brevican bands from **a**. Statistics by 2-way ANOVA with Tukey’s test for multiple comparisons. Band 1 likely corresponds to the full-length brevican, while band 2 likely corresponds to a truncated cleavage product.^121^ N = 6 mice per genotype. **c**. Raw TMT signal from proteomics for brevican. Statistics by 2-way ANOVA with Tukey’s test for multiple comparisons. N = 3 mice per genotype. **d**. Raw TMT signal from proteomics for hepaCAM. Statistics by 2-way ANOVA with Tukey’s test for multiple comparisons. N = 3 mice per genotype.

**Supplementary Figure 12.**
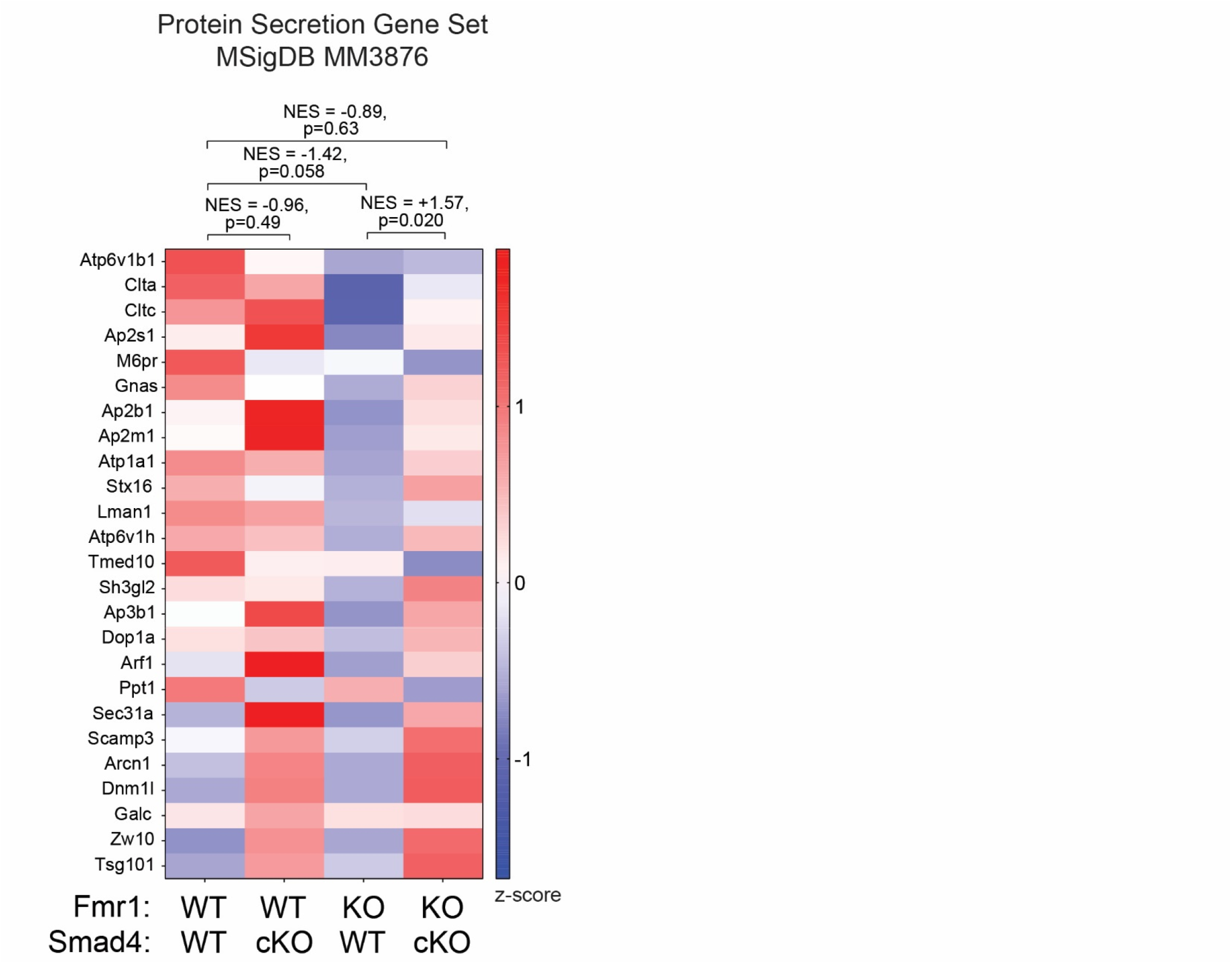
Smad4 cKO restores Fmr1 KO astrocyte deficit in secretory machinery. Z-scores of all detected proteins in the “protein secretion” gene set (MSigDB MM3876), averaged within each genotype.

